# Modulation of NF-κB signaling by *Alternaria* mycotoxins: *in vitro* and *in silico* insights into molecular mechanisms of immunosuppression in THP-1 monocytes

**DOI:** 10.64898/2026.07.06.736814

**Authors:** Vanessa Partsch, Francesco Crudo, Christian Schröder, Giorgia Del Favero, Doris Marko

## Abstract

*Alternaria* fungi produce various structurally diverse mycotoxins, several of which exhibit immunomodulatory properties. Among these, alternariol monomethyl ether (AME), alternariol (AOH), alterperylenol (ALTP), altertoxin I (ATX-I), and altersetin (AST) have been reported to suppress lipopolysaccharide (LPS)-induced inflammatory responses. However, the precise molecular mechanisms underlying these effects remain unclear.

The present study aimed to elucidate how these selected *Alternaria* mycotoxins (0.1-50 µM) target the NF-κB signaling pathway in THP-1 monocytes. Key components of the NF-κB cascade were analyzed by immunofluorescence microscopy, Western blotting and qRT-PCR. Nuclear translocation of NF-κB p65 and its phosphorylated form (p- NF-κB p65) was assessed by Western blot, while cytokine responses were determined at transcript (qRT-PCR) and protein (ELISA) levels. Moreover, *in silico* docking analyses were performed to investigate potential interactions of the toxins with IKKβ, and receptor-mediated crosstalk was studied using the glucocorticoid receptor (GR) antagonist RU486.

Co-treatment with RU486 attenuated the immunosuppressive effects of 1 and 5 µM AOH, indicating partial involvement of GR-dependent mechanisms. AME, AOH, ALTP, ATX-I, and AST increased total IκBα levels while reducing its phosphorylated form. Additionally, AST and ALTP decreased the protein levels of Toll-like receptor 4 (TLR4), the IκB kinase (IKK) complex, NF-κB p65, and p- NF-κB p65. While AOH (5 µM) and AST (25 µM) reduced nuclear translocation of p65 and p-p65, ALTP (2 µM) enhanced nuclear localization despite decreasing cytokine expression

Together, these findings suggest toxin-specific interference at multiple regulatory levels of NF-κB signaling and provide novel mechanistic insight into the immunomodulatory effects of *Alternaria* mycotoxins.

## 1. Introduction

*Alternaria* species are common phytopathogenic fungi that frequently contaminate a wide range of food and feed commodities, including cereals, oilseeds, fruits, and vegetables [1]. To date, more than 70 structurally distinct secondary metabolites have been identified, including compounds toxic to humans and animals collectively referred to as *Alternaria* mycotoxins. These toxins can be sub-categorized according to their chemical structures. Prominent examples include dibenzo-α-pyrones such as alternariol (AOH) and alternariol monomethyl ether (AME), perylene quinones like alterperylenol (ALTP) and altertoxin I (ATX-I), as well as structurally distinct compounds such as altersetin (AST) [1–3] (Fig 1). Although these toxins often co-occur in food and feed, toxicological evaluations have mainly focused on only a few single compounds like AOH and AME. Consequently, the biological activities and molecular targets of less well-studied toxins remain poorly understood. The lack of comprehensive toxicological data has also hindered the establishment of regulatory limits for these emerging mycotoxins.

**Figure 1:**
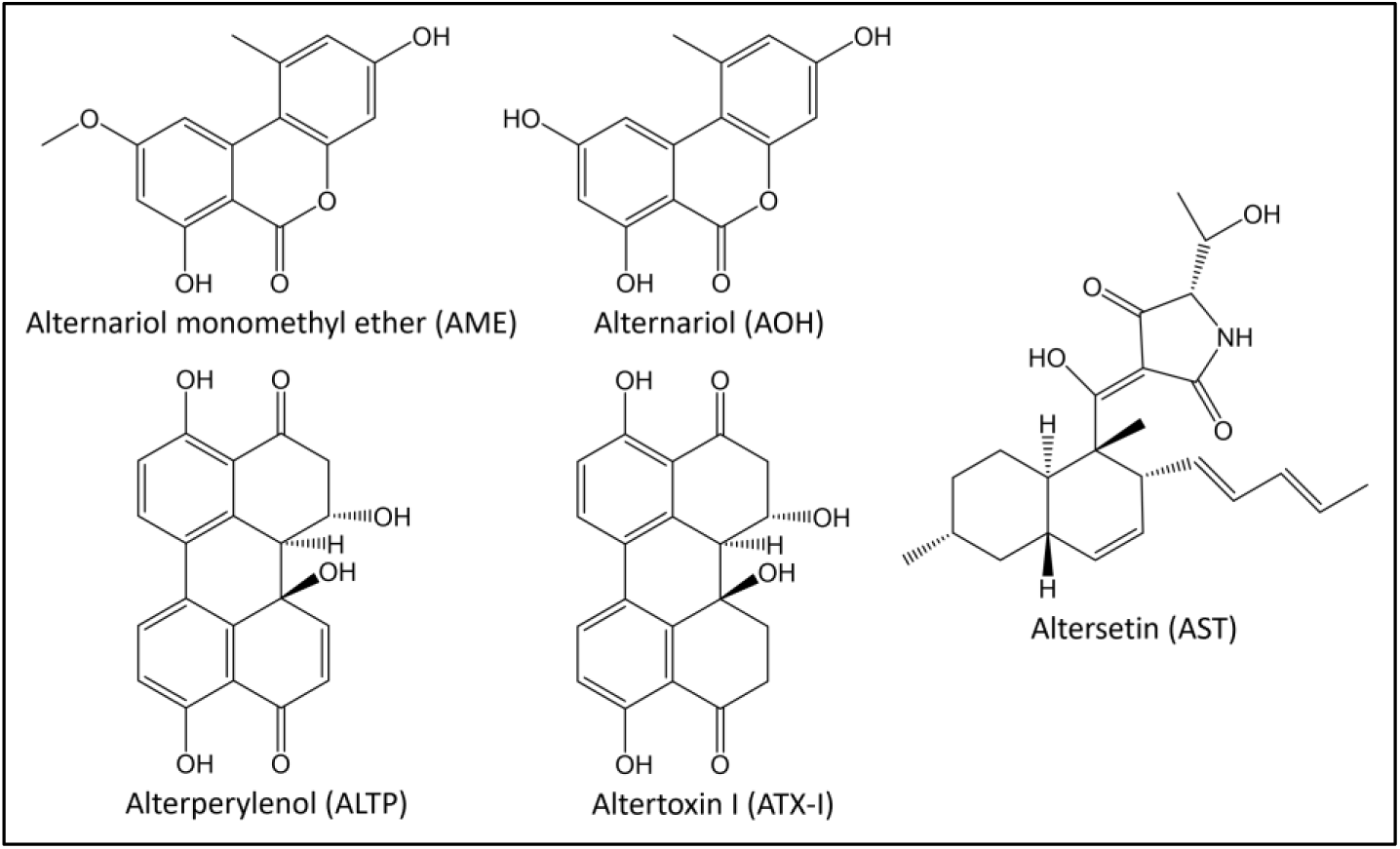
Chemical structures of the *Alternaria* mycotoxins alternariol monomethyl ether (AME), alternariol (AOH), alterperylenol (ALTP), altertoxin I (ATX-I) and altersetin (AST).

Several *Alternaria* mycotoxins have been reported to exert genotoxic, estrogenic, and immunomodulatory effects [4]. Among these, modulation of immune signaling is particularly important, as immunosuppression can increase susceptibility to infections or disrupt inflammatory homeostasis [5]. The innate immune system is a key target for such immunomodulatory compounds and is regulated, among others, by the nuclear factor kappa-light-chain-enhancer of activated B cells (NF-κB) pathway. NF-κB is a transcription factor activated downstream of pat-tern-recognition receptors, such as Toll-like receptor 4 (TLR4). These receptors respond to a wide range of ligands, including cytokines like interleukin (IL)-1β and tumor necrosis factor alpha (TNF-α), as well as microbial components like bacterial lipopolysaccharide (LPS). NF-κB comprises five related proteins that can form dimers, which in the canonical pathway are typically composed of p65 (RelA) and p50. In resting cells, these NF-κB dimers, are sequestered in the cytosol by inhibitory proteins belonging to the inhibitor of κB (IκB) family, primarily IκBα. Binding of ligands to the respective receptors leads to the activation of the IκB kinase (IKK) complex, consisting of IKKα, IKKβ, and the regulatory subunit NF-κB essential modulator (NEMO). Subsequently, the IKK phosphorylates the IκB proteins, marking them for ubiquitination and proteasomal degradation. As a result, the NF-κB dimers are released and translocate into the nucleus, where they promote the transcription of target genes. Many of these genes are involved in inflammation, immunity and cell survival. Notably, several pro-inflammatory cytokines such as IL-6, IL-8, IL-1β and TNF-α are under the control of this pathway [6].

Environmental and dietary xenobiotics have been shown to interfere with the NF-κB signaling cascade at different molecular levels. While some act as pro-inflammatory agents by inducing NF-κB activation, others suppress NF-κB activity and downstream cytokine release [7]. In recent years, several *Alternaria* mycotoxins were reported to inhibit NF-κB activation in immune cells as well as epithelial cell models. For instance, the mycotoxins AOH, AST, ALTP and ATX-I have been shown to potently inhibit the LPS-induced activation of the NF-κB signaling pathway in THP-1 monocytes [8–10]. Moreover, AOH and ALTP were found to downregulate several pro-inflammatory cytokines like IL-6 and IL-8 in the intestinal epithelial cell lines Caco-2 and HCEC-1CT, affecting both mRNA expression and protein secretion [11,12]. However, the underlying molecular mechanisms of the observed immunosuppressive effects are still to be elucidated.

Inflammation is regulated through multiple, interconnected control levels, with several intersecting signaling pathways contributing to its coordination. Besides the NF-κB pathway, the glucocorticoid receptor (GR) signalling is well known to play an important role in the regulation of the immune system. Upon binding glucocorticoids such as dexamethasone (Dex), the GR dissociates from its cytosolic chaperone complex, translocates into the nucleus, and modulates gene expression either by direct binding to glucocorticoid response elements (GREs) or through protein-protein interactions with other transcription factors [13]. Importantly, activated GR can physically interact with NF-κB subunits, a process known as transrepression, which interferes with NF-κB function and can alter both its nuclear translocation and DNA-binding capacity [14,15]. Whether a crosstalk between GR and NF-κB signaling contributes to the immunoinhibitory potential of *Alternaria* mycotoxins remains to be clarified.

Building upon this knowledge, the present study aimed to investigate the molecular mechanisms underlying the immunosuppressive effects of the *Alternaria* mycotoxins AOH, AME, ALTP, ATX-I, and AST on the NF-κB pathway. For this purpose, THP-1 Lucia^TM^ monocytes were co-exposed to LPS and the mycotoxins of interest. Given the complex nature of the NF-κB signaling pathway, changes in different constituents of the cascade were examined. To observe the behavior of multiple regulatory components, including TLR4, IκBα, IKK, and the NF-κB protein p65, the study employed immunofluorescence microscopy, Western blotting, and qRT-PCR. Moreover, *in silico* docking analyses were performed investigate potential interactions of the toxins with IKKβ. To assess whether the observed immunosuppressive effects are mediated through GR-dependent mechanisms, the NF-κB reporter gene assay was conducted in the presence of the GR antagonist RU486 (mifepristone). Finally, the impact of these mycotoxins on the pro- and anti-inflammatory cytokines IL-6, IL-8, IL-10 and TNF-α was evaluated using qRT-PCR and ELISA.

## 2. Materials and Methods

### 2.1. Materials

Cell culture media (Park Memorial Institute 1640 medium (RPMI 1640)), penicillin streptomycin (P/S) solution, fetal bovine serum (FBS), HEPES buffer and sodium pyruvate were purchased from Thermo Fisher Scientific Inc. (Waltham, USA). Zeocin® and Normocin® were obtained from Invivogen (San Diego, USA). Cell culture consumables were bought from Sarstedt AG + Co. KG (Nuembrecht, Germany), Carl Roth GmbH + Co. KG (Karlsruhe, Germany) and VWR International (Pennsylvania, USA).

LPS from *Escherichia coli*, Dex, phenylmethylsulfonyl fluoride (PMSF) and donkey serum were acquired from Sigma Aldrich (St. Louis, USA). The CellTiter Blue® (CTB) reagent was purchased from Promega Corp. (Fitchburg, USA), Mifepristone (RU486; ab120356) and Fluoroshield Mounting Medium with DAPI (ab104139) from Abcam (Cambridge, UK), adhesion slides (12 x 5 mm Ø) from Paul Marienfeld GmbH & Co. KG (Lauda-Königshofen, Germany), and dimethyl sulfoxide (DMSO), Triton-X 100, TAE buffer, and ROTI®Fix spray fixative from Carl Roth GmbH + Co. KG (Karlsruhe, Germany). PVDF membranes and filter papers for protein blotting were acquired from Bio Rad Laboratories (Hercules, USA), cOmplete™ and PhosSTOP™ and phosphatase inhibitor cocktails from Roche Holding AG (Basel, Switzerland), and the RIPA buffer from Merck KGaG (Darmstadt, Germany). The NE-PER™ Nuclear and Cytoplasmic Extraction Kit, the Halt™ Protease Inhibitor Cocktail, the Pierce™ BCA Protein Assay Kit, the Pierce^TM^ ECL Western Blotting Substrate, the High-Capacity cDNA Reverse Transcription Kit, the *Power* SYBR™ Green PCR Master Mix, PCR plates, ELISA Kits (human IL-6, IL8, IL-10 and TNF-α, uncoated) and the Stop Solution for TMB Substrates were purchased from Thermo Fisher Scientific Inc. (Waltham, USA).

Primary antibodies against IκBα (ab76429) were purchased from Abcam (Cambridge, UK), IκBα phospho-S32/S36 (A27385) from Antibodies.com LLC (St. Louis, USA), TLR4 (E5D8T), IKKα/β phospho-Ser176/180 (16A6), NF-κB p65 (D14E12), NF-κB p65 phospho-Ser536 (93H1), and GAPDH (D4C6R) from Cell Signaling Technology (Danvers, USA), α-Tubulin (sc-5286) from Santa Cruz Biotechnology Inc. (Dallas, USA), and Lamin B1 (33-2000) and the secondary antibody Alexa Fluor 568 donkey anti-rabbit (A10042) from Thermo Fisher Scientific Inc. (Waltham, USA). The anti-mouse IgG (7076S) and anti-rabbit IgG (7074S) horseradish peroxidase (HRP)-conjugated secondary antibodies were obtained from Cell Signaling Technology (Danvers, USA). The RNeasy Mini Kit and the primer for IκBα (Hs_NFKBIA_1_SG; QT00014266), IKKα (Hs_NFKBIA_1_SG; QT00014266), IL-6 (Hs_IL6_1_SG; QT00083720), IL-8 (Hs_CXCL8_1_SG; QT00000322), IL-10 (Hs_IL10_1_SG; QT00041685) and TNF-α (Hs_TNF_1_SG; QT00029162) were obtained from Qiagen N.V. (Hilden, Germany), while the primers for NF-κB p65 (reverse: TCAGCCTCATAGAA-GCCATC, forward: GCACAGATACCACCAAGACC), TLR4 (reverse: GCTTATCTGAAGGTGTTGCACAT, forward: CAGAG-TTTCCTGCAATGGATCA), and GAPDH (reverse: GATTTGGTCGTATTGGGCGC, forward: TTCCCGTTCTCAGCCTTGAC) were acquired from Eurofins Scientific SE (Luxembourg City, Luxembourg). The *Alternaria* mycotoxin AME was acquired from Biomol GmbH (Hamburg, Germany), AOH from Sigma Aldrich (St. Louis, USA), ALTP from Cfm Oskar Tropitzsch (Marktredwitz, Germany), ATX-I from Cayman Chemicals Company (Ellsworth, USA), and AST from Molport (Riga, Latvia).

### 2.2. Cell culture

THP-1 Lucia™ monocytes (InvivoGen, San Diego, USA) were maintained in RPMI 1640 medium supplemented with 10% FBS, 25 mM HEPES, 1% P/S (100 U/ml), and 100 µg/ml Normocin®. To maintain reporter gene stability, 100 µg/ml Zeocin® was added every second passage. Cells were cultured at 37 °C in a humidified atmosphere containing 5% CO₂ and passaged twice per week. Mycoplasma contamination was routinely monitored.

### 2.3. Mycotoxin treatment and dosage information

For all experiments, THP-1 Lucia™ monocytes were pre-incubated for 2 h with different concentrations of the *Alternaria* mycotoxins AOH, AME, ATX-I, ALTP and AST, a solvent control (0.25% DMSO), and a negative control (1 µM Dex). Subsequently, NF-κB activation was induced by adding 10 ng/ml LPS to all wells except the solvent control, followed by an additional incubation period of 18 h or in the case of qRT-PCR and ELISA for further 3 h and 18 h. LPS-treated cells (10 ng/ml) served as the positive control for NF-κB activation. Table 1 provides an overview of the *Alternaria* mycotoxins and the respective concentrations tested in each assay. For the mRNA and protein analyses only non-cytotoxic concentrations were selected.

**Table 1:** Concentrations of the *Alternaria* mycotoxins tested in the respective assays.

| | Tested concentrations [μM] | | | | | | |
| --- | --- | --- | --- | --- | --- | --- | --- |
| | NF-κB assay | CTB assay | IF microscopy | Western blot (complete) | Western blot (cytosolic/nuclear extracts)** | qRT-PCR | ELISA |
| AME* | 0.1-20 | 0.1-20 | 0.1, 5, 10 | - | - | - | - |
| AOH | 0.1-20 | 0.1-20 | 0.1, 1, 5, 10 | 0.1, 1, 5 | 5 | 0.1, 1, 5 | 0.1, 1, 5 |
| ALTP | 0.1-20 | 0.1-20 | 0.1, 1, 2 | 0.1, 1, 2 | 2 | 0.1, 1, 2 | 0.1, 1, 2 |
| ATX-I | 0.1-20 | 0.1-20 | 0.1, 2, 20, 50 | - | - | - | - |
| AST | 0.1-20 | 0.1-20 | 0.1, 1, 10, 25 | 1, 10, 25 | 25 | 1, 10, 25 | 1, 10, 25 |

\*alternariol monomethyl ether (AME), alternariol (AOH), alterperyleneol (ALTP), altertoxin I (ATX-I), altersetin (AST)
\*\*cytosolic and nuclear proteins were separated to investigate the nuclear translocation of NF-κB p65 and p-NF-κB p65

### 2.4. NF-κB reporter gene assay

To assess the potential involvement of GR signaling in mycotoxin-mediated immunosuppression, the NF-κB reporter gene assay was performed in the presence of the glucocorticoid receptor antagonist RU486. THP-1 monocytes were seeded into 96-well plates at a density of 0.1 × 10⁶ cells/well and were pre-incubated with 1 µM RU486 for 1 h. Subsequently, exposure of the cells to the *Alternaria* mycotoxins (0.1-20 µM) and the different controls were performed as described in section 2.3. All conditions were tested in the presence and absence of RU486 to enable direct comparison. After a total incubation time of 20 h, plates were centrifuged at 140 rcf for 2 min, and 20 µl of the supernatants were transferred to a white 96-well plate. NF-κB activity was measured according to the manufacturer’s instructions by measuring luciferase activity after adding 80 µl of the Quanti-Luc™ reagent. Luminescence was detected with a Cytation 3 microplate reader (BioTek Instruments, Winooski, USA) using Gen5 software (version 3.08).

### 2.5. CellTiter-Blue® (CTB) assay

The CTB assay was performed to confirm that the effects observed in the toxicological assays were not artifacts caused by cytotoxicity. The experimental conditions were identical to those described for the NF-κB reporter gene assay (see section 2.4). At the end of the incubation, the CTB reagent (1:10 dilution in supernatant) was added to each well, followed by 2 h exposure. Cells treated with 0.01% Triton X-100 served as a positive control for cytotoxicity. Plates were centrifuged at 140 rcf for 2 min, and 100 µL of the supernatants were transferred to a black 96-well plate. Fluorescence was measured at 560/590 nm (λ_ex_/λ_em_) with a microplate reader.

### 2.6. Immunofluorescence (IF) microscopy

To assess the impact of the *Alternaria* mycotoxins AME (0.1-10 µM), AOH (0.1-10 µM), ALTP (0.1-2 µM), ATX-I (0.1-50 µM) and AST (0.1-25 µM) on the protein levels of the inhibitory protein IκBα and its phosphorylated form (p-IκBα), IF microscopy was performed. THP-1 monocytes were seeded into 96-well plates at a density of 0.1 × 10⁶ cells/well and incubation conditions were applied as described in section 2.3. At the end of the incubation period, cells were fixed on adhesion slides. In detail, 35 µl of the cell suspension was transferred onto one spot of the slide, followed by 30 min incubation to allow attachment of the cells. The supernatant was removed and cells were fixed using ROTI®Fix spray fixative, followed by storage at -20 °C overnight. Cells were permeabilized with ice-cold methanol, and blocking was performed with 2% donkey serum in PBS (2.9 mM KH₂PO₄, 5.4 mM KCl, 273.8 mM NaCl, 30.9 mM Na₂HPO₄·2H₂O; in distilled water; pH 7.2) for 1 h at room temperature as previously described [16]. Subsequently, cells were incubated with the primary antibody (1:100 diluted in 2% donkey serum) for 1 h at room temperature. After removing the antibody solution, cells were washed three times with PBS for 15 min each. Incubation with the corresponding secondary antibody (1:400 in 2% donkey serum) was performed for 1 h at room temperature, followed by another three washing steps with PBS for 15 min each. Finally, mounting medium containing DAPI was applied to counterstain cell nuclei. Slides were sealed to preserve the staining and were stored at 4 °C until imaging. Images of nuclei, IκBα, and p-IκBα were acquired using a Zeiss LSM 800 (Carl Zeiss Microscopy, Oberkochen, Germany), equipped with a Plan Apochromat 63×/1.4 oil objective (zoom 1.5). At least three images from different optical fields were captured per spot. Single-cell analysis was performed with a minimum of 21 cells per biological replicate. Image analysis was conducted using the Fiji software (ImageJ 1.54f).

### 2.7. Western blot

#### 2.7.1. Analysis of the whole protein extract

Changes in protein levels of p-IKKα/β, TLR4, NF-κB p65, and p-NF-κB p65 were assessed in THP-1 monocytes exposed to the *Alternaria* mycotoxins AOH (0.1-5 µM), ALTP (0.1-2 µM) and AST (1-25 µM) using Western blot analysis. Since the immunosuppressive effects of these toxins were more pronounced and apparent at lower concentrations, it was decided not to conduct further analyses using ATX-I. Furthermore, AME was excluded due to the co-occurrence of cytotoxicity (see Supplementary Figure S1).

For Western blot cells were seeded in 12-well plates at a density of 1 × 10⁶ cells/well and treated according to section 2.3. After incubation, three technical replicates were pooled to obtain a higher cell number. Cells were centrifuged, washed with ice-cold PBS, and lysed on ice in RIPA buffer supplemented with 1 mM PMSF, Complete® protease inhibitor, and PhosSTOP® phosphatase inhibitor cocktails according to manufacturer’s protocol. Lysates were sonicated for 5 min and centrifuged at 18,000 rcf for 20 min at 4 °C. Protein concentrations in the resulting supernatants were determined using the Pierce BCA Protein Assay Kit according to manufacturer’s instructions. Equal amounts of protein (20 µg per lane, diluted in RIPA and loading buffer) were loaded onto 4% polyacrylamide stacking gels and separated on 10% polyacrylamide separation gels at 90 V for 110 min. Proteins were transferred onto PVDF membranes by wet blotting (400 mA, 110 min). Membranes were blocked in 5% fat-free milk in Tris-buffered saline containing 0.1% Tween-20 (TBS-T) for 1 h at room temperature. Incubation with the specific primary antibodies against p-IKKα/β, TLR4, NF-κB p65, p-NF-κB p65 and GAPDH as a housekeeping control (1:1000 in 3% BSA in TBS-T) was performed overnight at 4 °C. After washing three times with TBS-T (8 min each), membranes were incubated for 1 h at room temperature with HRP-conjugated secondary antibodies (anti-mouse IgG and anti-rabbit IgG; 1:10000 in 1% BSA in TBS-T). After three additional washes with TBS-T, protein bands were visualized using the Pierce ECL Western Blotting Substrate and imaged with the LAS-4000 Image Reader (Fujifilm, Minato, Japan). Band intensities were quantified using the MultiGauge software (version 3.0).

#### 2.7.2. Analysis of cytosolic and nuclear protein extracts

To assess potential changes in the nuclear translocation of NF-κB and its phosphorylated form, proteins of THP-1 monocytes were fractionated into nuclear and cytosolic extracts. Cells were seeded in T-25 flasks at a density of 10 × 10⁶ cells/flask and treated with the highest concentrations of AOH (5 µM), ALTP (2 µM) and AST (25 µM) as described in Section2.3. Following incubation, cells were collected and proteins were fractionated using the NE-PER™ Nuclear and Cytoplasmic Extraction Kit, complemented with Halt™ protease inhibitor and PhosSTOP® phosphatase inhibitor cocktails according to the manufacturer’s instructions. Protein concentrations were determined using the Pierce BCA Protein Assay Kit and equal amounts of protein (10 µg per lane) from each fraction were loaded onto gels and transferred to PVDF membranes. Lamin B1 and α-tubulin were used as nuclear and cytosolic markers respectively to confirm fraction purity. Both antibodies were diluted 1:250 in 3% BSA in TBS-T. The remaining procedure of the Western blot was performed according to the protocol described for total protein extracts in section 2.7.1.

### 2.8. Quantitative real time PCR (qRT-PCR)

To determine the effects of AOH (0.1-5 µM), ALTP (0.1-2 µM) and AST (1-25 µM) on the mRNA transcript levels of NF-κB p65, IκBα, IKK, TLR4, and the cytokines IL-6, IL-8, IL-10 and TNF-α, two-step qRT-PCR was performed. THP-1 monocytes were seeded in 12-well plates at a density of 1 × 10⁶ cells/well and treated as described in section 2.3. Following incubation, supernatants were collected and stored at -80 °C for further protein analyses (see section 2.9). To obtain a higher cell number, two technical replicates were pooled for RNA extraction. Cells were washed with ice-cold PBS, and total RNA was extracted using the RNeasy Mini Kit, according to manufacturer’s instructions. RNA concentration and purity were assessed using a NanoDrop 2000 spectrophotometer (Thermo Fisher Scientific, Waltham, USA). For complementary DNA (cDNA) synthesis, 1 µg of total RNA per sample was reverse transcribed using the High-Capacity cDNA Reverse Transcription Kit following the manufacturer’s protocol. Gene-specific cDNA was then amplified by qRT-PCR with a QuantStudio 3 PCR System (Thermo Fisher Scientific, Waltham, USA). Reactions were carried out using *Power* SYBR™ Green PCR Master Mix, along with gene-specific primer assays in 96-well plates with a final reaction volume of 20 µL per well. The amplification protocol consisted of an initial activation step at 95 °C for 15 min, followed by 40 cycles of denaturation at 94 °C for 15 sec, annealing at 55 °C for 30 sec, and extension at 70 °C for 30 sec. A final melting curve analysis was performed to confirm amplification specificity (1 min at 65 °C, heating in 0.5 °C increments to 94 °C, holding at 95 °C for 15 sec). Relative mRNA expression levels were calculated using the 2^−ΔΔCT^ method. CT values of target genes were normalized to the average CT values of the housekeeping gene GAPDH and fold changes were determined relative to the corresponding samples of the positive control.

### 2.9. Enzyme-linked immunosorbent assay (ELISA)

To assess whether the *Alternaria* mycotoxins AOH (0.1-5 µM), ALTP (0.1-2 µM) and AST (1-25 µM) affect cytokine secretion, supernatants collected after cell treatment (see section 2.8) were analyzed by ELISA. Protein levels of IL-6, IL-8, IL-10, and TNF-α were quantified using commercial ELISA kits according to the manufacturer’s instructions. Absorbance was measured at 450 nm using a microplate reader

### 2.10. *In silico* analysis

To investigate potential interactions of the five *Alternaria* mycotoxins with IKKβ, molecular docking experiments were performed. The selection of IKKβ for *in silico* analyses was based on its central role in the canonical NF-κB signaling pathway, where its catalytic activity is required for the phosphorylation and subsequent degradation of IκB proteins [17]. This choice was further supported by the experimental findings of the present study, which indicated interference with upstream regulatory events consistent with impaired IKK activity. In addition, previous studies have reported that AOH can interact with the ATP-binding pocket of several kinases, supporting the hypothesis that *Alternaria* mycotoxins may directly target kinase activity [18,19].

The crystal structure of human IKKβ (PDB ID: 4KIK) was obtained from the Protein Data Bank [20]. The co-crystallized ligand and water molecules were removed, and only one monomer of the dimeric structure was used for docking to keep the computational effort reasonable. Docking simulations were performed with AME, AOH, ALTP, ATX-I and AST using SwissDock implementing the Attracting Cavities 2.0 algorithm [21,22]. Predicted binding poses were grouped into clusters and ranked according to the estimated binding free energy (ΔG), which represents an empirical approximation used to prioritize ligand conformations rather than an absolute thermodynamic value [21]. For each predicted cluster, the conformation with the lowest ΔG value was selected for further analysis. These selected poses were subsequently analyzed using Visual Molecular Dynamics (VMD) with respect to their localization within the kinase structure and their potential interactions with functionally relevant regions, including the ATP-binding site [23].

### 2.11. Statistical analyses

Statistical analyses were performed using Origin Pro® 2022 software (OriginLab, Northampton, USA). All experiments were conducted in technical triplicates or technical duplicates (qRT-PCR and ELISA) and in at least three biological replicates. Data were normalized to the positive control and are presented as mean + standard deviation (SD) of the biological replicates. Statistical significance between the positive control and individual treatments was determined using Student’s *t*-test. Differences among multiple concentrations of a single mycotoxin were analyzed by one-way ANOVA followed by Fisher-LSD post-hoc test. For IF, statistical comparisons between different concentrations were performed using the Kruskal-Wallis test, followed by Dunn’s multiple comparison test for post-hoc analysis.

## 3. Results

### 3.1. Impact of the GR on the immunosuppressive and cytotoxic effects of *Alternaria* mycotoxins

The involvement of the GR in the immunosuppressive effects of selected *Alternaria* mycotoxins (AME, AOH, ALTP, ATX-I, and AST) was investigated using an NF-κB reporter gene assay in the presence of 1µM of the GR antagonist RU486. THP-1 Lucia™ monocytes were co-exposed to five concentrations of each mycotoxin (0.1-20 µM) and 10 ng/ml LPS to induce NF-κB signaling. NF-κB activity was subsequently compared between incubations performed in the presence and absence of RU486. As shown in Figure 2A, AOH significantly suppressed LPS-induced NF-κB activation starting at 1 µM. Co-treatment with the GR inhibitor RU486 counteracted the immunosuppressive effects of AOH, as no reduction in NF-κB activity was observed at 1 µM. Moreover, at 5 µM AOH, NF-κB signaling was significantly increased in the presence of RU486 compared to treatment with AOH alone (p<0.05). In contrast, no significant differences between exposures performed with or without RU486 were observed for any of the other mycotoxins (Supplementary Figure 1A-D).

**Figure 2:**
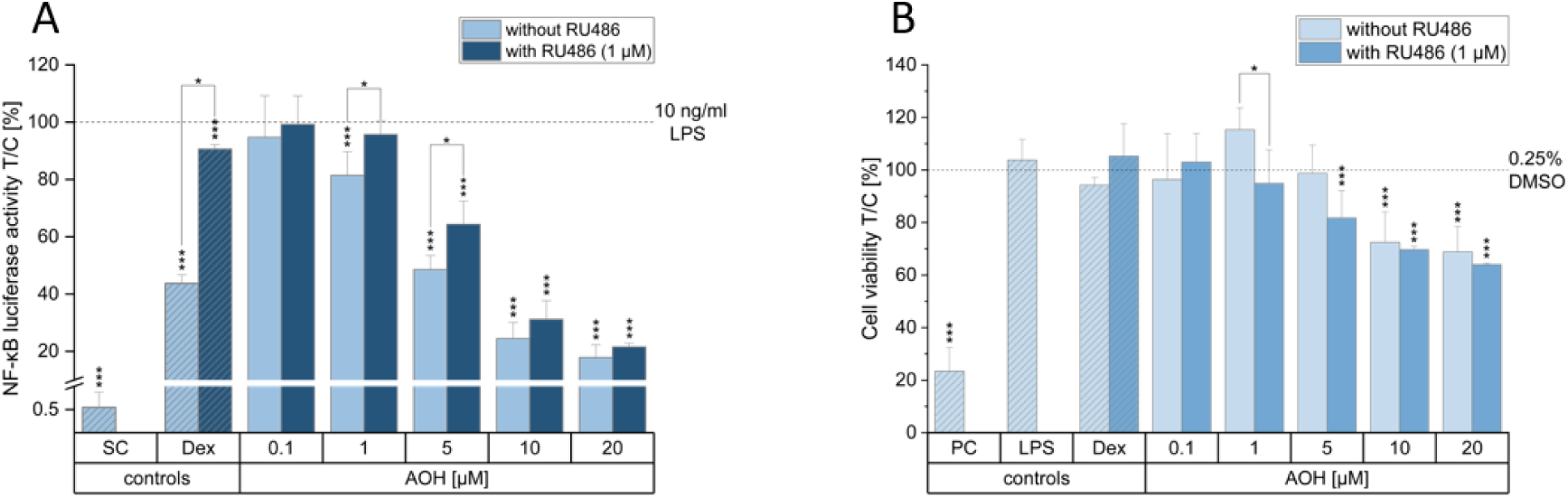
Immunoinhibitory and cytotoxic effects of alternariol (AOH) in the presence and absence of the glucocorticoid receptor inhibitor RU486. THP-1 Lucia™ monocytes were either pre-incubated with 1 µM RU486 for 1 h or exposed to AOH (0.1-20 µM) alone for 2 h, followed by co-stimulation with 10 ng/ml lipopolysaccharide (LPS) for further 18 h. Dexamethasone (Dex; 1 µM) served as negative control, 0.25% DMSO as solvent control (SC), and in panel (B) 0.01% Triton X-100 as a positive control (PC) for cytotoxicity. Immunosuppressive effects were assessed by the NF-κB reporter gene assay (A) and cytotoxicity by the CellTiter-Blue® (CTB) assay (B). Data are presented as mean + SD of at least three independent experiments and are expressed relative to the positive control (10 ng/ml LPS) in (A) and relative to the SC in (B), indicated by dotted lines. Statistical significance compared to the positive control and between treatments with and without RU486 was evaluated using Student’s t-test (*p<0.05, **p<0.01, and ***p<0.001).

Cytotoxicity was assessed in parallel using the CTB assay. As shown in Figure 2B AOH induced a significant reduction in cell viability starting at 10 µM (p<0.05). Co-treatment with RU486 and 5 µM AOH also resulted in minor but statistically significant cytotoxic effects. However, cell viability remained above 80%. The respective cytotoxicity data for the remaining four *Alternaria* mycotoxins are provided in the supplementary materials (Supplementary Figure 1E-H).

### 3.2. Impact of *Alternaria* mycotoxins on the IκBα and p-IκBα protein levels

To investigate possible changes in the cytosolic protein content of the inhibitory protein IκBα and p-IκBα upon exposure to the immunosuppressive *Alternaria* mycotoxins AME, AOH, ALTP, ATX-I and AST, IF microscopy analyses were conducted. As shown in Figure 3, all tested mycotoxins induced a concentration-dependent upregulation of IκB accompanied by a downregulation of p-IκBα. The only exception was ATX-I, which did not affect cytosolic p-IκBα levels. Nevertheless, exposure to 2 µM ATX-I resulted in a significant upregulation of IκBα (Figure 3E) as compared to the to the positive control (10 ng/ml LPS). AOH, ALTP, and AST caused increased IκBα levels starting at 1 µM. Interestingly, at 10 µM AOH IκBα levels declined again and did not significantly differ from those of the positive control (Figure 3C). In contrast, p-IκBα levels were already reduced by ALTP and AST at 0.1 µM, while for AOH the lowest effective concentration was 1 µM (Figure 3C,D,F). With regards to AME, both an upregulation of IκBα and a downregulation of p-IκBα were already observed at 0.1 µM (Figure 3B). Corresponding representative fluorescence images of all controls and of the mycotoxin concentrations inducing the strongest effects are depicted in Figure 3A.

**Figure 3:**
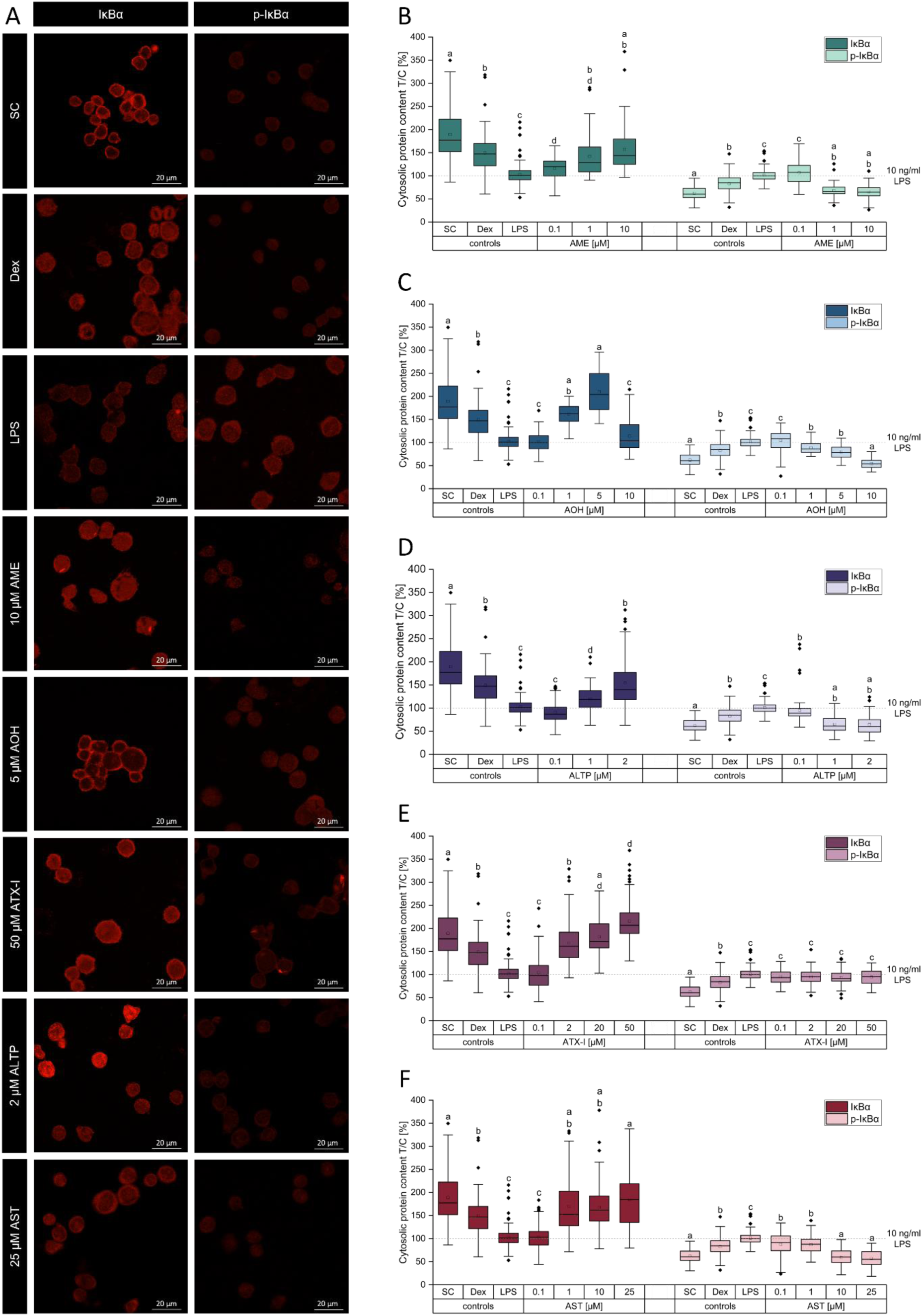
Effects of the *Alternaria* mycotoxins alternariol monomethyl ether (AME), alternariol (AOH), alterperylenol (ALTP), altertoxin I (ATX-I), and altersetin (AST) on cytosolic IκBα and phosphorylated IκBα (p-IκBα) levels. THP-1 monocytes were exposed to the *Alternaria* mycotoxins for 2 h followed by co-stimulation with 10 ng/ml lipopolysaccharide (LPS) for further 18 h. Dexamethasone (Dex; 1 µM) served as a negative control, while 0.25% DMSO was used as solvent control (SC). Cytosolic IκBα and p-IκBα contents were assessed by immunofluorescence microscopy. Results are presented relative to the positive control (10 ng/ml LPS), indicated by dotted lines. Representative confocal microscopy images of the controls (SC, 1µM Dex, 10 µg/ml LPS) and of the mycotoxin concentrations showing the greatest differences compared to the positive control (10 µM AME, 5 µM AOH, 2 µM ALTP, 50 µM ATX-I, and 25 µM AST) are shown in panel (A). For all conditions, three independent experiments were performed, and ≥21 cells were selected for quantification per experiment Statistical significance between treatments and the positive control (LPS) was evaluated using the Kruskal–Wallis test followed by Dunn’s multiple-comparison test. Groups that do not share a common letter (a–d) differ significantly at p < 0.05, with all treatments compared directly to the LPS-stimulated control.

### 3.3. Impact of AOH, ALTP and AST on TLR4, p-IKKα/β, NF-κB p65 and p-NF-κB p65 protein levels

To further elucidate the impact of AOH, ALTP, and AST on key components of the TLR4-NF-κB signaling pathway, including TLR4, p-IKKα/β, NF-κB p65, and p-NF-κB p65, protein expression levels were analyzed by Western blotting. Analyses were restricted to these three *Alternaria* mycotoxins, as they emerged as the most active compounds in this study. Quantitative protein expression data together with representative corresponding bands of the Western blots are shown in Figure 4. Exposure of the cells to ALTP and AST lead to a downregulation of all four analyzed proteins. Specifically, AST significantly downregulated TLR4 (p < 0.001) and p-NF-κB p65 (p<0.001) only at 25 µM (Figure 4A,D) whereas reductions in p-IKKα/β (p < 0.05) and NF-κB p65 (p < 0.01) were observed already at concentrations ≥ 10 µM (Figure 3B, C). In contrast, ALTP significantly reduced protein levels of TLR4 (p<0.001) and NF-κB p65 (p<0.01) only at 2 µM (Figure 4 A, C), while p-IKKα/β (p<0.01) and p-NF-κB p65 (p<0.001) were decreased starting at 1 µM (Figure 3B, D). On the contrary, AOH only caused a slight upregulation of p-NFκB p65 protein levels with significant differences compared to the positive control already at 0.1 µM (p<0.01; Figure 4D).

**Figure 4:**
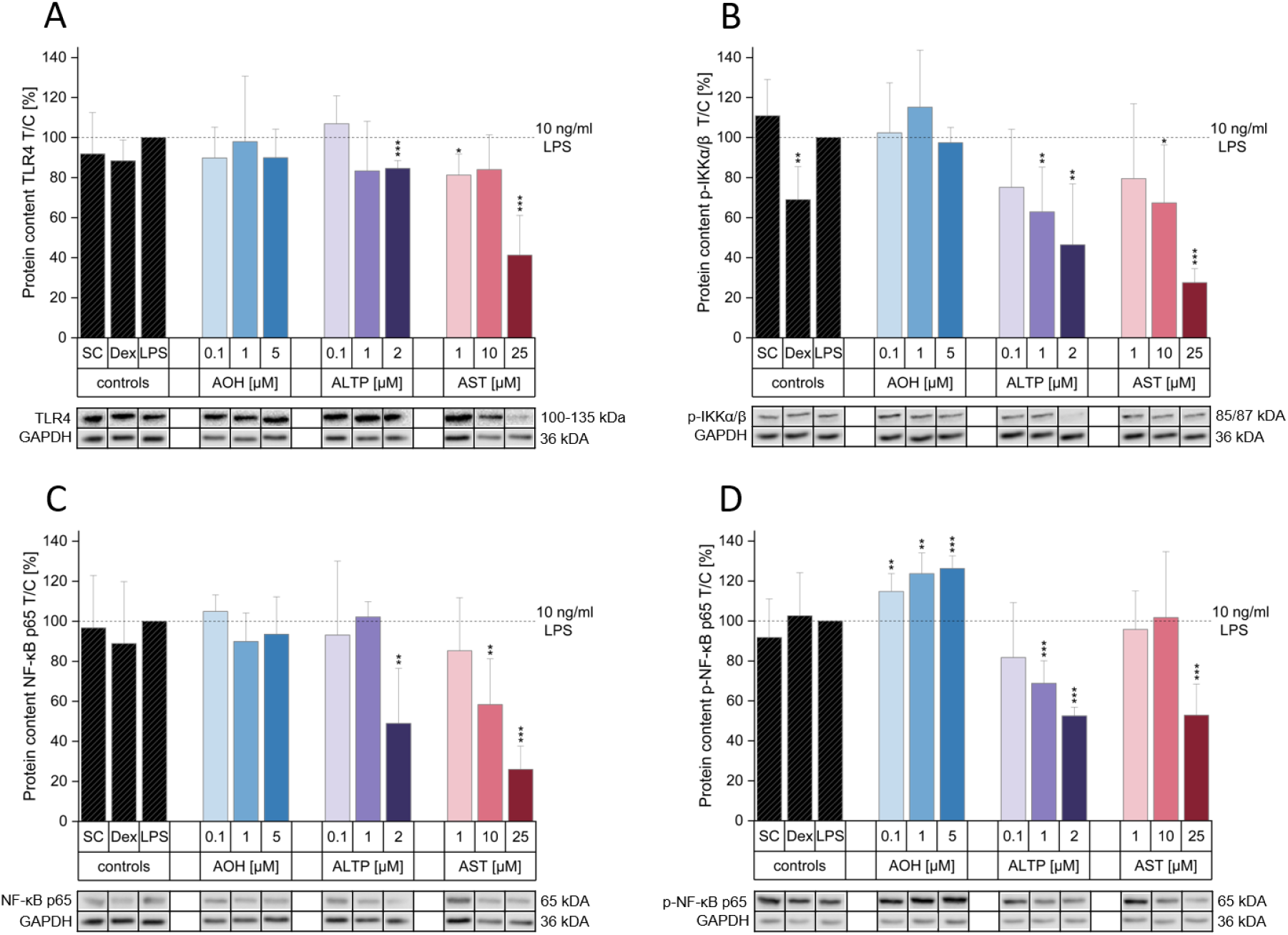
Effects of alternariol (AOH), alterperylenol (ALTP), and altersetin (AST) on key proteins of the NF-κB signaling pathway. THP-1 monocytes were exposed to the *Alternaria* mycotoxins for 2 h followed by co-stimulation with 10 ng/ml lipopolysaccharide (LPS) for further 18 h. Dexamethasone (Dex; 1 µM) served as a negative control, while 0.25% DMSO was used as solvent control (SC). Protein expression of Toll-like receptor 4 (TLR4, A), phosphorylated IκB kinase α/β (p-IKKα/β, B), NF-κB p65 (C), and phosphorylated NF-κB p65 (p-NF-κB p65, D) was assessed by Western blot analysis. GAPDH was used as a housekeeping protein for normalization. Data are presented as mean + SD of at least three independent experiments and are expressed relative to the positive control (10 ng/ml LPS), indicated by dotted lines. Statistical significance compared to the positive control was evaluated using Student’s *t*-test (*p<0.05, **p<0.01, and ***p<0.001). Bands shown below the respective graphs were derived from a single blot and were cut and rearranged, as indicated by dividing lines between lanes, to improve visualization.

### 3.4. Changes in nuclear translocation of NF-κB p65 and p-NF-κB p65

To assess possible changes in the nuclear translocation of NF-κB p65 and p-NF-κB p65, Western blot analyses were performed using cytosolic and nuclear protein extracts. The highest tested concentrations of AOH, ALTP, and AST, namely 5 µM, 2 µM, and 25 µM respectively, were selected for this analysis. Figure 5 depicts the distribution of the proteins in cytosolic and nuclear fractions, normalized to total protein levels, as well as the corresponding ratios relative to the positive control (10 ng/mL LPS). Exposure to AOH and AST resulted in reduced nuclear translocation of both NF-κB p65 and p-NF-κB p65. In contrast, ALTP treatment led to an increased relative nuclear-to-cytosolic distribution compared to the positive control for both proteins. However, absolute nuclear protein levels were reduced for all three mycotoxins.

**Figure 5:**
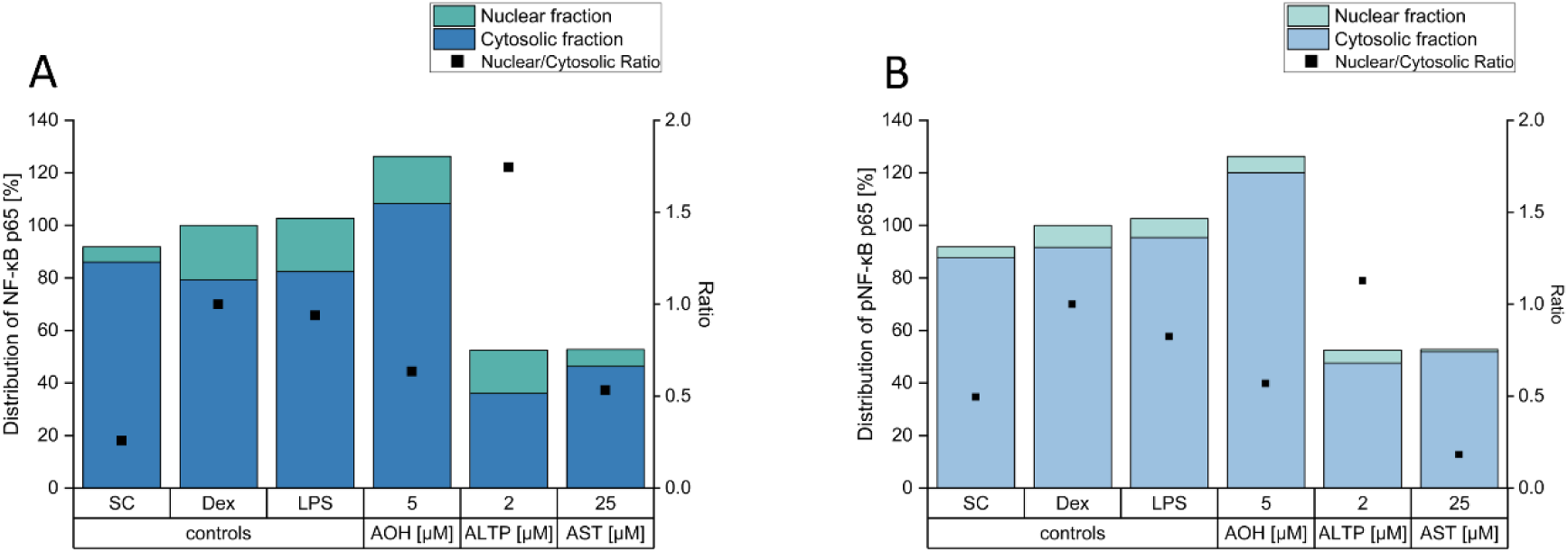
Effects of alternariol (AOH), alterperylenol (ALTP), and altersetin (AST) on cytosolic and nuclear NF-κB p65 and phosphorylated NF-κB p65 (p-NF-κB p65) protein levels. THP-1 monocytes were exposed to the *Alternaria* mycotoxins for 2 h followed by co-stimulation with 10 ng/ml lipopolysaccharide (LPS) for further 18 h. Dexamethasone (Dex; 1 µM) served as a negative control, while 0.25% DMSO was used as a solvent control (SC). Cytosolic and nuclear fractions were prepared, and protein expression levels of NF-κB p65 (A) and p-NF-κB p65 (B) were assessed by Western blot analysis. α-Tubulin and Lamin B1 were used for normalization of cytosolic and nuclear fractions, respectively. Data were collected from at least three independent experiments, and the ratios are expressed relative to the positive control (10 ng/ml LPS).

### 3.5. Impact of AOH, ALTP and AST on TLR4, IKK, IκBα and NF-κB p65 mRNA levels

To assess potential changes in the transcriptional regulation of key NF-κB pathway components and to complement the protein expression and translocation analyses, qRT-PCR experiments were performed. Changes in the mRNA levels of TLR4, IKK, IκBα, and NF-κB p65 in THP-1 monocytes after exposure to AOH, ALTP, and AST for 5 h and 20 h were elucidated. As shown in Figure 6A and E, AST caused minor decreases in TLR4 mRNA levels after both 5 h and 20 h at all tested concentrations, whereas AOH and ALTP remained largely ineffective. For IKK, exposure to ALTP for 5 h resulted in a slight upregulation of transcription starting at 1 µM, while after 20 h a decrease in IKK mRNA levels was observed (Figure 6B,F). The most pronounced effects were detected for IκBα, where 2 µM ALTP reduced transcript levels to below 0.5-fold after both 5 h (p<0.05) and 20 h (p<0.001) of exposure (Figure 6C,G). No consistent changes were observed for NF-κB p65 transcription. However, exposure to 25 µM AST resulted in a slight but significant downregulation of NF-κB p65 mRNA levels after 20 h (p<0.001) (Figure 6D,H).

**Figure 6:**
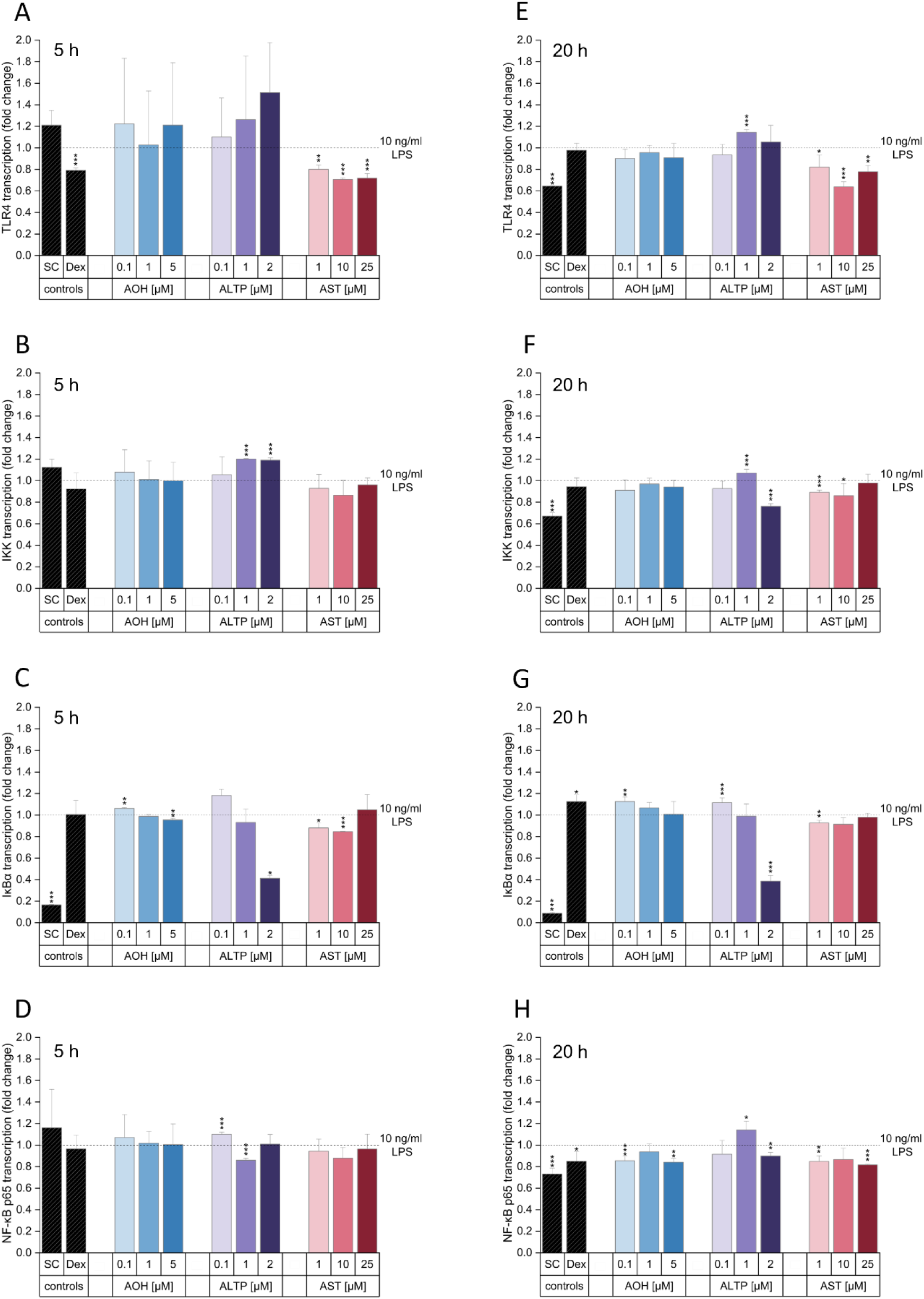
Impact of the *Alternaria* mycotoxins alternariol (AOH), alterperylenol (ALTP), and altersetin (AST) on the relative gene transcription of TLR4, IKK, IκBα, and NF-κB p65. THP-1 monocytes were exposed to the *Alternaria* mycotoxins for 2 h and stimulated with 10 ng/ml lipopolysaccharide (LPS) for further 3 h (A-D) or 18 h (E-H). Dexamethasone (Dex; 1 µM) served as a negative control, while 0.25% DMSO was used as solvent control (SC). Gene expression changes were examined by qRT-PCR, and results were calculated as relative gene transcription (2^−ΔΔCT^), normalized to GAPDH and compared to the positive control (10 ng/ml LPS), indicated by dotted lines. Data are presented as mean + SD from at least three independent experiments. Statistical significance between mycotoxin-treated samples and the positive control was evaluated using Student’s *t*-test (*p<0.05, **p<0.01, and ***p<0.001).

### 3.6. Integration of mRNA and protein expression data of key NF-κB pathway components

To provide a comprehensive overview of the results obtained from the mRNA and protein expression experiments, the data are integrated and presented in Figure 7. Only AOH, ALTP, and AST were included, as a complete and consistent dataset across all experimental endpoints was generated for these compounds. Accordingly, Figure 7 summarizes their impact on key components of the NF-κB signaling pathway as described in sections 3.2-3.5, thereby enabling direct comparison across endpoints and facilitating data interpretation. Specifically, Figure 7A presents a simplified schematic overview of the pathway, whereas Figure 7B illustrates the observed changes in protein and mRNA levels as well as alterations in nuclear protein content. In contrast, ATX-I was not considered further due to its less pronounced immunosuppressive effects at lower concentrations, while AME was excluded because of the co-occurrence of cytotoxicity (see Supplementary Figure S1).

**Figure 7:**
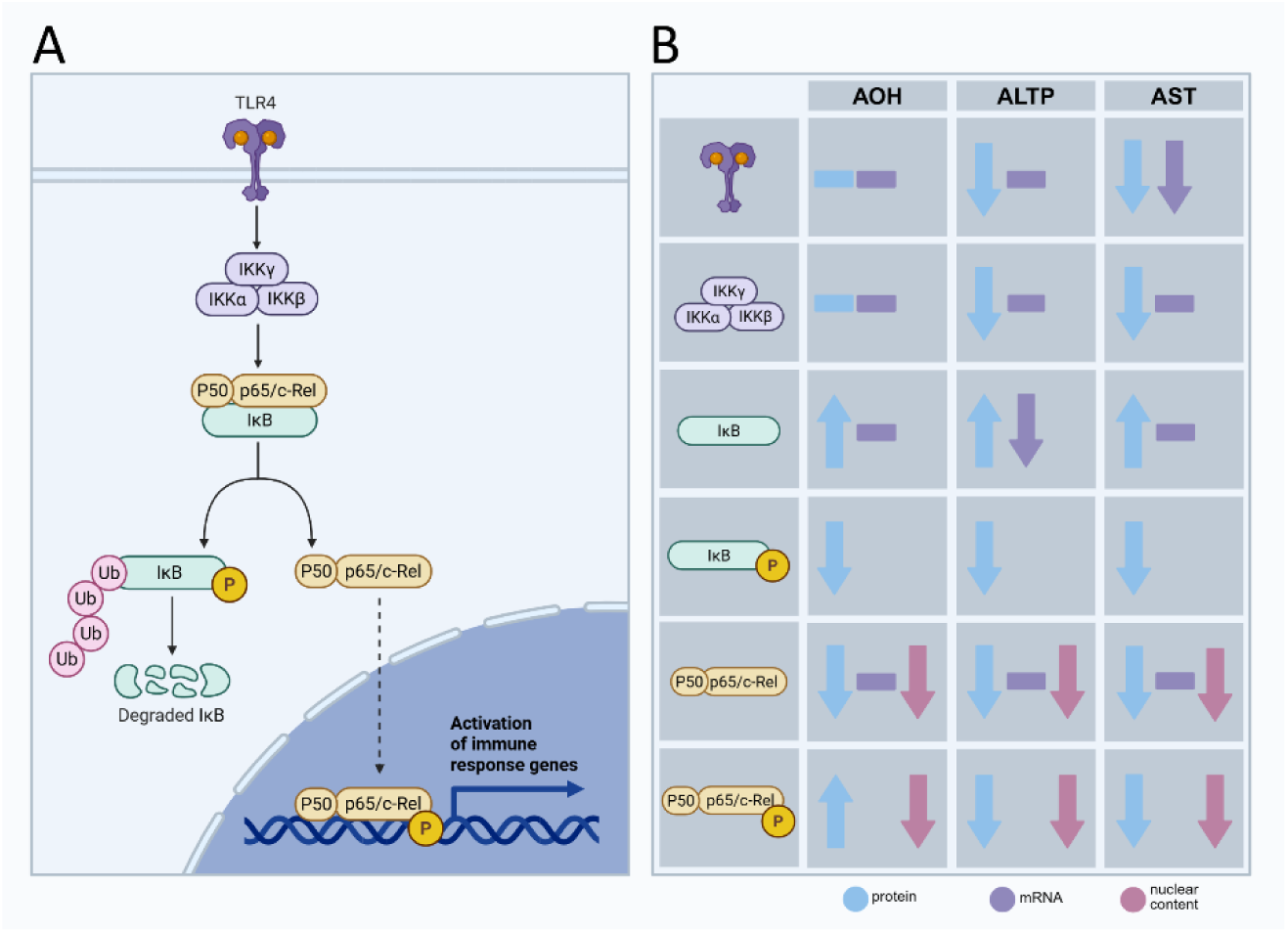
Overview of the NF-κB signaling pathway (A) and impact of the *Alternaria* mycotoxins alternariol (AOH), alterperylenol (ALTP) and altersetin (AST) on key components of the signaling cascade compared to the positive control (10 ng/ml LPS) (B). Upregulation is indicated by upward arrows, downregulations by downward arrows, and the absence of an effect by a hyphen.

### 3.7. *In silico* docking analysis of AME; AOH, ALTP, ATX-I and AST with IKKβ

To investigate potential interactions of the five Alternaria mycotoxins with IKKβ, molecular docking experiments were performed, as this kinase represents a key upstream regulator of the canonical NF-κB pathway and a potential target consistent with the observed experimental effects. Figure 8 shows a representative structure of the IKKβ dimer used for the *in silico* analyses, along with all identified docking positions of the five mycotoxins, ranked according to predicted binding free energy (ΔG, SwissDock scoring). Close-up views illustrate the conformation of each respective mycotoxin in its most favorable binding position. For kinase inhibition of IKKβ through ligand binding to the ATP-binding pocket, the amino acids Met96, Glu97, Tyr98, Cys99, Asp166, Leu167, and Gly168 are most commonly described as being involved in ligand interactions and are highlighted in red in Figure 8A [24]. Analysis of the potential interactions of the different toxins with IKKβ revealed that the most favorable conformations of AME and AOH were located within the ATP-binding pocket, as indicated by their proximity to all seven of the mentioned amino acids. Only three and five less likely additional binding sites, respectively, were identified, suggesting a specific interaction of these two dibenzo-α-pyrones with the ATP-binding site (Figure 8B,C). For the perylene quinones ALTP and ATX-I, a total of eight and ten possible interaction sites were identified, respectively. Only few of the most favorable conformations were located within the ATP-binding pocket, indicating rather unspecific interactions between these mycotoxins and the kinase subunit (Figure 8D,E). In contrast, AST did not show any favorable interaction within the ATP-binding pocket, but two out of five possible confirmations were clearly favored by the toxin (Figure 8F).

**Figure 8:**
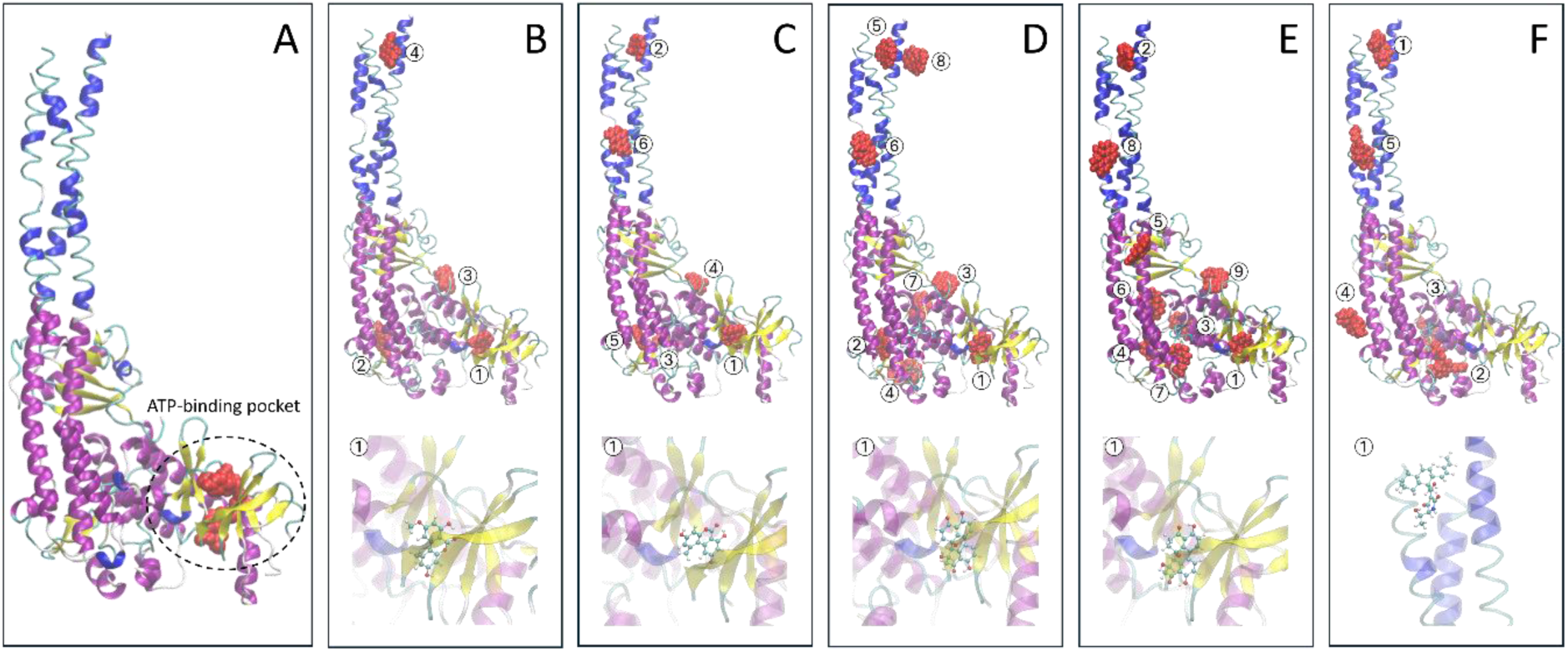
Representative structure of the IκB kinase β (IKKβ) dimer used for *in silico* docking analyses (A) and predicted binding sites of the *Alternaria* mycotoxins alternariol monomethyl ether (AME, B), alternariol (AOH, C), alterperylenol (ALTP, D), altertoxin I (ATXI, E), and altersetin (AST, F). Panel A shows the ATP-binding pocket, with key amino acid residues involved in ligand binding highlighted in red. Panels B–F display all identified docking poses for each toxin, ranked according to their predicted probability of interaction. Close-up views illustrate the conformation and orientation of each mycotoxin in its most favorable binding position within the kinase structure.

Detailed information on all identified docking clusters, including their assignment to the corresponding binding positions shown in Figure 8B–F, the number of conformations per cluster, and the ΔG values of the conformationswith the lowest predicted binding free energy within each cluster, is provided in the Supplementary Materials (Supplementary Tables 1–5).

### 3.8. Impact of AOH, ALTP and AST on IL-6, IL-8, IL-10 and TNF-α mRNA and protein levels

To further assess downstream inflammatory responses, the transcriptional and protein expression levels of the NF-κB-regulated cytokines IL-6, IL-8, IL-10, and TNF-α were analyzed. Specifically, mRNA levels were quantified by qRT-PCR, whereas cytokine secretion was determined using ELISA following exposure of THP-1 monocytes to AOH, ALTP, and AST for 5 h and 20 h. IL-10 protein levels were below the limit of detection under all tested conditions and could therefore not be quantified by the applied method (data not shown).

While THP-1 cells were largely unresponsive to the applied AOH concentrations after 5 h at both the mRNA and protein level, exposure for 20 h resulted in an upregulation of IL-6 mRNA and a downregulation of IL-8 mRNA at 5 µM AOH (Figure 9B). These transcriptional changes were reflected at the protein level and were accompanied by a slight but significant increase in TNF-α secretion for all tested AOH concentrations compared to the positive control (10 ng/ml LPS; Figure 9D).

**Figure 9:**
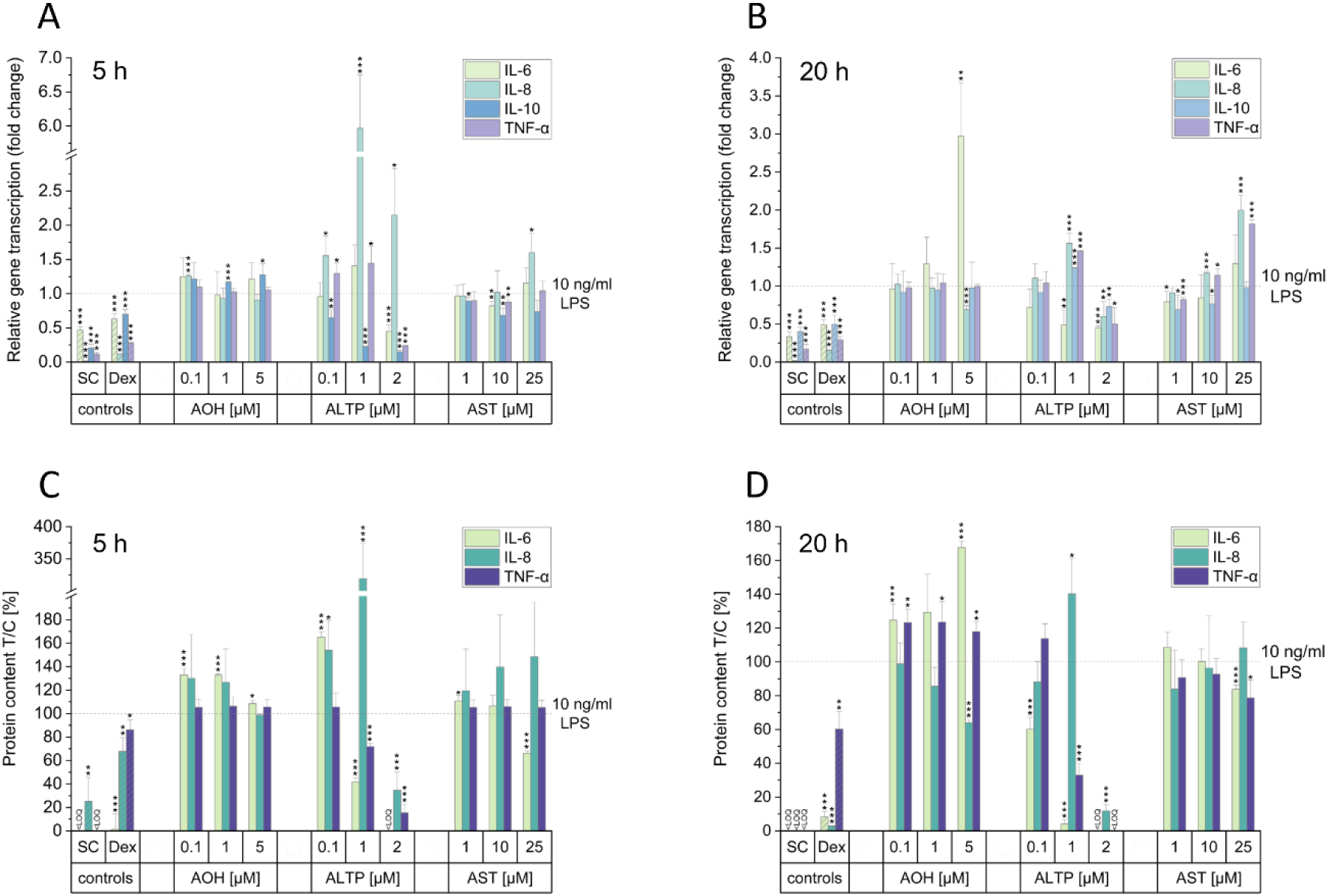
Impact of the *Alternaria* mycotoxins alternariol (AOH), alterperylenol (ALTP), and altersetin (AST) on the relative gene transcription and cytokine secretion levels of IL-6, IL-8, IL-10 and TNF-α. THP-1 monocytes were exposed to the *Alternaria* mycotoxins for 2 h and stimulated with 10 ng/ml lipopolysaccharide (LPS) for further 3 h (A, C) or 18 h (B,D). Dexamethasone (Dex; 1 µM) served as a negative control, while 0.25% DMSO was used as solvent control (SC). Gene expression changes were examined by qRT-PCR (A,B), and results were calculated as relative gene transcription (2^−ΔΔCT^), normalized to GAPDH. Cytokine secretion levels were assessed by ELISA (C,D). Data are presented as mean + SD from at least three independent experiments and are expressed relative to the positive control (10 ng/ml LPS), indicated by dotted lines. Statistical significance between mycotoxin-treated samples and the positive control was evaluated using Student’s *t*-test (*p<0.05, **p<0.01, and ***p<0.001).

For ALTP, a more complex pattern was observed since it induced downregulation of IL-10 mRNA starting at 0.1 µM, and of IL-6 and TNF-α mRNA only at 2 µM. In contrast, IL-8 transcription was strongly upregulated, with a six-fold increase at 1 µM, which decreased at 2 µM (Figure 9A). These gene expression changes were largely mirrored at the protein level, since IL-6 concentrations fell below the LOQ and TNF-α secretion was reduced to 20% at 2 µM ALTP. Consistent with the mRNA data, IL-8 protein levels were highly elevated at 1 µM (∼300%) and decreased to around 40% at 2 µM (Figure 9C). After 20 h of ALTP exposure, responses were less pronounced. In detail IL-6 mRNA levels were reduced starting at 1 µM, while IL-8, IL-10, and TNF-α mRNA levels were slightly increased at 1 µM but decreased at 2 µM (Figure 9B). On the protein level, IL-6 secretion was already reduced starting at 0.1 µM, and TNF-α secretion decreased at 1 µM, with both falling below the LOQ at 2 µM. Similar to the transcriptional analysis, IL-8 protein levels increased at 1 µM and then fell below 20% at 2 µM (Figure 9D).

After 5 h of exposure to AST, only a minor upregulation of IL-8 mRNA levels was observed at 25 µM, however, this effect was not reflected at the protein level. Instead, a decrease in IL-6 secretion was detected at 25 µM (Figure 9A,C). In contrast, 20 h of incubation resulted in increased transcription levels of IL-8 and TNF-α starting at 10 µM. These transcriptional effects were not mirrored at the protein level, where only minor decreases in IL-6 and TNF-α secretion were observed (Figure 9B,D).

## 4. Discussion

The group of *Alternaria* mycotoxins includes over 70 chemically diverse compounds. Some of these molecules are known to exert a wide range of adverse effects and can be found in a variety of food and feed. Among other toxicological endpoints, it was previously described that some these compounds possess immunomodulatory properties [8–10]. However, the precise molecular mechanisms underlying the actions of these *Alternaria* mycotoxins are yet still unknown. Thus, the present study aimed to elucidate the specific targets of these toxins within the NF-κB signaling pathway, thereby providing insights into their mechanistic actions at the molecular level.

Glucocorticoids represent one of the most potent endogenous anti-inflammatory regulators and exert their effects primarily through activation of the GR. Since GR signaling is a well-established negative regulator of NF-κB-mediated inflammatory responses, its involvement in the immunosuppressive effects of *Alternaria* mycotoxins was investigated. Among other mechanisms, GR activation can suppress cytokine gene expression by interfering with transcription factors such as NF-κB. These mechanisms are complex and include direct protein-protein interactions with NF-κB subunits, stabilization or induction of IκBα, and GR binding to specific DNA elements, resulting in transrepression of NF-κB-dependent gene transcription [14,15]. AOH has previously been described as an immunosuppressive compound [8–10], which is in line with the findings of the present study. Consistent with earlier reports, AOH significantly suppressed LPS-induced NF-κB activity in THP-1 monocytes (Figure 2A). Importantly, co-exposure of cells to AOH and the GR antagonist RU486 attenuated the immunoinhibitory effects of the mycotoxin at 1 µM and 5 µM. These findings suggest that GR signaling contributes to AOH-mediated suppression of NF-κB activity. However, a more in-depth investigation is required to elucidate the exact GR-mediated immunoinhibitory mechanisms. Notably, the immunosuppressive activity of the other four tested mycotoxins was not affected by co-incubation with the GR antagonist (Supplementary Figure 1), indicating that these compounds likely exert their effects via GR-independent modes of action.

Investigations of key components of the NF-κB pathway upon exposure of the cells revealed distinct modulation patterns in response to different mycotoxins. AOH did not have any impact on the protein levels nor the mRNA levels of TLR4 and IKK (Figure 4A,B; Figure 6A,B,E,F). These results partially align with observations by Rønning et al. (2019), who reported no changes in TLR4 transcript levels in THP-1 macrophages exposed to urolithin A, a bacterial metabolite structurally related to AOH. Interestingly, Del Favero et al. (2020b) [25] demonstrated that AOH can increase membrane fluidity and alter the interplay between caveolin-1 (Cav-1) and TLR4. Cav-1, a structural protein of caveolae, plays a crucial role in membrane organization as well as receptor distribution and turnover [26]. Moreover, it can act as a scaffold for TLR4 and modulate the recruitment of downstream adaptor proteins such as MyD88, thereby fine-tuning NF-κB activation [27]. Consequently, AOH-induced modulations in membrane organization may contribute to the observed suppression of NF-κB activity, independent of total receptor abundance.

However, alterations could be measured at the level of cytosolic IκBα regulation. Specifically, starting at 1 µM, corresponding to the lowest concentration at which immunosuppressive effects were detected, AOH induced an upregulation of total IκBα protein alongside a decrease in p-IκBα (Figure 3C). Under non-stimulated conditions, IκB proteins inhibit NF-κB activity by retaining NF-κB dimers in the cytoplasm and preventing nuclear translocation. Increased IκBα levels therefore promote an inactive state and reduce NF-κB-dependent transcription. In contrast, phosphorylation of IκBα triggers its ubiquitination and proteasomal degradation, releasing NF-κB dimers for phosphorylation, nuclear translocation, and activation of target gene transcription [6]. The changes in IκBα and p-IκBα levels upon the exposure to AOH therefore provide a mechanistic explanation for the observed suppression of NF-κB reporter activity. In line with these findings, Komatsu et al. (2018) [28] reported that urolithin A increased IκBα protein levels in LPS-stimulated RAW 264.7 macrophages, supporting the hypothesis that structurally related dibenzo-α-pyrones may suppress NF-κB signaling by enhancing IκB-dependent cytoplasmic retention of NF-κB. Although phosphorylation levels of IKKα/β were not altered by AOH (Figure 4B), the pronounced reduction in phosphorylation of IκBα together with an increase in total IκBα hints towards impaired functional IKK activity. The IKK complex consists of the catalytic subunits IKKα and IKKβ and non-catalytic regulatory subunit NEMO, with IKKβ being the primary kinase component of the IKK complex phosphorylating IκBα in the canonical NF-κB pathway [6,29]. Using an *in silico* approach, Aichinger et al. (2020) [18] identified casein kinase II (CK2) as a potential target of AOH. This observation was later complemented by Crudo et al. (2024) [19] who suggested inhibitory interactions of AOH with kinases involved in the formation of γH2AX. In these studies, the toxin was found to favorably dock within the ATP-binding pocket of CK2 as well as the serine-protein kinases ATM, DNA-PKcs, and the serine/threonine-protein kinase ATR and persist therein [18,19]. Since ATP-binding sites in kinases are typically highly conserved [30], there is a reasonable possibility that AOH may not only target CK2 and kinases involved in γH2AX formation, but may also act as a rather non-specific kinase inhibitor. Based on these findings, the present study assessed the affinity of AOH for the ATP-binding site of IKKβ using *in silico* docking analysis. The results indicated that the most favorable conformations of AOH were located within the ATP-binding pocket (Figure 8C). Given that CK2 is involved in the co-regulation of NF-κB activation [31], and the potential for AOH to interact with kinase ATP-binding sites, it appears conceivable that AOH may modulate kinase activity within the NF-κB signaling cascade. Consequently, this may contribute to the reduced phosphorylation of IκBα and the subsequent suppression of NF-κB signaling observed in the present study.

Consistently, the present data show a reduced nuclear translocation of both NF-κB p65 and p-NF-κB p65 following exposure of THP-1 monocytes to 5 µM AOH (Figure 5). These findings are in line with Groestlinger et al. (2022) [32] who reported reduced nuclear translocation of NF-κB p65 in HCEC-1CT cells co-exposed to 10 µM AOH and IL-1β. Similarly, urolithin A has been reported to suppress NF-κB signaling by downregulating the translocation of NF-κB p65 to the nucleus and inhibiting its binding to κB-responsive DNA elements in LPS-stimulated RAW 264.7 macrophages [28,33]. In the present study, AOH induced a slight but significant increase in p-NF-κB p65 levels, while total NF-κB p65 protein remained unchanged (Figure 4CD). Importantly, this slight increase in phosphorylation did not translate into enhanced NF-κB activity. In contrast, urolithin A has been shown to reduce both total and p-NF-κB p65 levels in THP-1 macrophages, highlighting compound- and cell-type-specific differences in the regulation of NF-κB signaling [34]. Together, these findings suggest that AOH might interfere with nuclear translocation and/or DNA binding, thereby resulting in overall immunosuppression.

Similar considerations apply to the interpretation of the results obtained for AME. Although the monomethyl ether of AOH induced a comparable upregulation of IκBα and downregulation of its phosphorylated form starting at 1 µM (Figure 3B) cytotoxic effects were observed at the same concentration (Supplementary Figure 1E). Furthermore, the *in silico* docking analysis indicated that the most favorable conformations of AME were located within the ATP-binding pocket of IKKβ (Figure 8B). Together with the findings of Aichinger et al. (2020) [18], who reported binding of AME to the ATP-binding site of CK2 and proposed a potential to affect kinase activity, these results suggest that AME may interact with kinase ATP-binding sites. However, given the observed overlap between pathway modulation and cytotoxicity, as also reported by Partsch et al. (2026) [12] the immunosuppressive potential of AME remains controversial, and it was not included in investigations of the transcriptional or protein level. Thus, further in-depth studies are needed to clarify its molecular effects.

On the contrary, AST was recently identified as a novel immunosuppressive *Alternaria* mycotoxin targeting the NF-κB signaling pathway in THP-1 monocytes and macrophages [10,12]. In the present study, AST induced a distinct response pattern compared to AOH. At the receptor level, 25 µM AST led to a reduction of TLR4 protein (Figure 4A). Along the downstream signaling cascade, a more homogeneous suppression was observed, including reduced levels of p-IKKα/β (Figure 4B), p-IκBα (Figure 3F), NF-κB p65 and p-NF-κB p65 (Figure 4C,D), as well as decreased nuclear translocation of NF-κB p65 and p-NF-κB p65 (Figure 5A,B). In accordance with this inhibitory pattern, cytosolic IκBα levels were increased (Figure 3F). These findings are consistent with Kuang et al. (2022) [35] who reported that proliferatins A-C, structurally related fungal metabolites, reduce phosphorylation of IKKα/β and NF-κB p65 and impair nuclear translocation of p65 in macrophages. While proliferatins did not alter TLR4 protein levels, their overall inhibitory effects on upstream NF-κB signaling support the hypothesis that structurally related compounds such as AST may interfere with early regulatory steps of the pathway. This is consistent with the observed downregulation of transcript and protein levels of TLR4 in the present study (Figure 4A, Figure 6A, E), as receptor expression and activation can be reduced by specific TLR4 inhibitors [36]. Consequently, this may result in a uniform downregulation of multiple key components along the NF-κB cascade. In line with this, a slight but significant downregulation of NF-κB p65 mRNA was observed (Figure 6H) which was also reflected on the protein level (Figure 4C). *In silico* docking analysis did not identify a clear preference of AST for the ATP-binding pocket of IKKβ, with the most probable poses located at alternative regions of the kinase (Figure 8F). Accordingly, no conclusions regarding a direct interaction of AST with the ATP-binding pocket or modulation of IKKβ activity can be drawn from these data. Overall, AST induced a broad attenuation of NF-κB signaling across multiple regulatory levels of the pathway. This suggests interference with upstream signaling events or a general attenuation of signaling capacity. However, the primary molecular target remains to be elucidated.

The highly potent immunoinhibitory perylene quinone ALTP, as reported by Crudo et al. (2025) [9] and Partsch et al. (2026) [12], was also found to modulate proteins of the NF-κB signaling in THP-1 monocytes, albeit with a distinct pattern compared to AOH and AST. Along the signaling cascade, ALTP exposure led to downregulation of p-IKKα/β (Figure 4B) and p-IκBα (Figure 3D), while cytosolic IκBα protein levels were increased (Figure 3D). Moreover, a downregulation of the IkBα mRNA was observed after 5 h and 20 h incubation (Figure 6C,G). The discrepancy between mRNA and protein levels may be explained by post-transcriptional regulatory mechanisms and altered protein stability and turn-over. Since IκBα degradation depends on IKK-mediated phosphorylation followed by ubiquitination and proteasomal degradation, the observed reduction in p-IκBα likely leads to stabilization of the protein despite reduced mRNA expression. This would promote cytosolic retention of NF-κB and contribute to the attenuation of downstream signaling [6]. Consistent with these *in vitro* observations, the *in silico* docking analysis revealed that ALTP exhibited multiple potential binding conformations on IKKβ, though only a subset of these were located within the ATP-binding pocket (Figure 8D). In line with these results, Aichinger et al. (2020) [18] reported that the structural analogue ATX-II favorably docked to the ATP-binding site of CK2 and inhibited enzyme activity, albeit with lower affinity than AOH. Together, these findings suggest that ALTP may be able to interfere with IKKβ, possibly leading to a reduced phosphorylation of IκBα. Interestingly, in contrast to AOH and AST, nuclear translocation of NF-κB p65 and its phosphorylated form appeared elevated compared to the positive control which was reflected by an elevated nuclear-to-cytosolic ratio (Figure 5). This occurred despite reduced total and phosphorylated NF-κB p65 protein levels (Figure 4C,D) and decreased NF-κB reporter activity (Supplementary Figure 1B). The apparent discrepancy points towards a relative nuclear enrichment, rather than true increases in nuclear protein abundance. Accordingly, increased nuclear localization did not translate into functional NF-κB activation, suggesting inhibition at the level of downstream transcriptional regulation. The regulation of immune responses involves extensive crosstalk between signaling pathways, including modulation by the aryl hydrocarbon receptor (AhR). Beyond its role as a sensor of xenobiotics and environmental stimuli, AhR has been shown to interact with NF-κB signaling at multiple levels, and to exert context-dependent immunomodulatory effects [37]. In particular, AhR can form complexes with RelA (i.e. NF-κB p65), resulting in suppression of NF-κB-dependent transcription [38]. In a study carried out by Hohenbichler et al. (2020) [39] the potential of several *Alternaria* mycotoxins to induce Cytochrome P450 (CYP) 1 activity, a classical downstream target of AhR signaling, was investigated. Since the structurally related perylene quinones ATX-I and ATX-II, were postulated to activate the AhR pathway, it appears plausible that ALTP may similarly function as an AhR agonist. Thus, AhR activation may represent a potential explanation for reduced NF-κB reporter activity despite increased relative nuclear localization of p- NF-κB p65. Taken together, ALTP may exert its immunosuppressive effects through transcriptional interference rather than solely through upstream inhibition of the NF-κB pathway.

ATX-I has previously been described as an immunosuppressive compound in LPS-stimulated THP-1 monocytes and macrophages [9,12], which is consistent with the present findings (Supplementary Figure 1C). Analysis of IκBα by immunofluorescence microscopy revealed a concentration-depended upregulation of the cytosolic protein content, while p-IκBα remained unaffected (Figure 3E). In contrast to the other *Alternaria* mycotoxins these effects were only observed at higher concentrations and were less pronounced. Therefore, no further transcriptional or protein analyses were performed for ATX-I. *In silico* docking analysis revealed a rather unspecific interaction pattern of ATX-I with IKKβ. Among the ten predicted binding conformations, only a few were located within the ATP-binding pocket, while several others were distributed across alternative regions of the kinase (Figure 8E). This indicates no clear preference of the toxin for the catalytic site. Together with the findings for ALTP and previously reported data for the structurally related ATX-II [18], these results suggest that perylene quinones may interfere with kinase activity, although likely with lower specificity than AOH. Interestingly, Del Favero et al. (2020a) [16] reported that ATX-II did not alter IκBα levels, but reduced NF-κB p65 nuclear translocation. The authors postulated a possible crosstalk between reactive oxygen species (ROS) signaling and NF-κB regulation. In this context, ATX-II and AOH were previously shown to trigger the release of the transcription factor nuclear factor erythroid-derived 2-like 2 (Nrf2) from its binding protein KEAP-1 and increase its nuclear translocation [32,40,41]. Activated Nrf2 can bind to IKKβ and suppress NF-κB signaling [42], providing a potential explanation for the immunosuppressive effects observed for AOH, ATX-II, and possibly ALTP. However, so far comparable evidence is lacking for ATX-I.

Given that NF-κB is a central regulator of pro- and anti-inflammatory cytokine expression, the observed upstream effects are expected to translate into alterations in the transcription and secretion of NF-κB dependent cytokines such as IL-6, IL-8, IL-10 and TNF-α [43,44]. However, the cytokine responses observed in the present study were highly toxin-specific and did not always directly reflect the modulation of upstream NF-κB signaling. For AOH, no major changes were observed after 5 h exposure (Figure 9A,C). However, following 20 h exposure, IL-6 mRNA and secretion were upregulated, whereas IL-8 levels decreased (Figure 9B,D). These findings only partially align with previous reports, as AOH has been described to decrease IL-6 and IL-8 transcription and secretion in various cell lines [8,11,12,32,45]. Similarly, while Kollarova et al. (2018) [8] reported reduced TNF-α and increased IL-10 secretion, the present study showed increased TNF-α secretion (Figure 9D) and elevated IL-10 mRNA (Figure 9A). However, a comparable increase in TNF-α secretion was reported by Partsch et al. (2026) [12] in Caco-2 cells. Notably, IL-10 protein levels were below the detection limit of the applied ELISA method and therefore reliable secretion data for this cytokine cannot be provided. MicroRNAs (miRNAs) are known to be involved in the regulation of immune signaling. The observed increase in IL-6 and TNF-α may be explained by previously described modulation of miR-155 and miR-146a by AOH [8], as these miRNAs are important regulators of NF-κB-dependent cytokine expression [46–49]. For AST, increased IL-8 and TNF-α mRNA levels were not reflected at the protein level, where IL-6 and TNF-α secretion were reduced (Figure 9). The observed discrepancy between mRNA expression and protein release suggests post-transcriptional or translational regulation. ALTP exhibited a concentration-dependent, biphasic effect. At the highest concentration tested (2 µM), all cytokine levels were downregulated, whereas at 1 µM, IL-8 was strongly upregulated before declining at 2 µM (Figure 9). Consistent with the present results, ALTP was previously shown to downregulate cytokine expression at both the mRNA and protein levels in HCEC-1CT and Caco-2 cells [12]. The transient upregulation of IL-8 at lower concentrations may indicate involvement of additional regulatory pathways, particularly the AhR. It can form transcriptionally active dimers with the NF-κB family member RelB, promoting IL-8 expression [50,51]. In contrast, the AhR can also interact with signal transducer and activator of transcription 1 (STAT1), inhibiting NF-κB transcriptional activity at promoter regions leading to a reduced IL-6 expression [52]. Furthermore, the association of activated AhR with the NF-κB p65 subunit has been shown to result in the inhibition of TNF-α production. In addition, AhR activation can induce negative regulators of NF-κB, such as the suppressor of cytokine signaling (SOCS2), which further modulate TNF-α expression [53]. These mechanisms may explain the concentration-dependent cytokine modulation observed for ALTP. Data on cytokine modulation by AST remain limited. However, supporting the present findings, Kuang et al. (2022) [35] showed a downregulation of IL-6, IL-8 and TNF-α in LPS-stimulated RAW264.7 macrophages treated with proliferatins A-C, hypothesized to be regulated through a TLR4-dependent mechanism. Of note, Partsch et al. [12] reported no significant changes in cytokine levels in Caco-2 or HCEC-1CT cells following AST exposure. Taken together, these findings indicate that downstream cytokine responses may not solely be regulated by NF-κB activation, but are more likely fine-tuned by multiple regulatory layers, including feedback loops, crosstalk with other signaling pathways, exposure duration, and cell-type specific factors. Consequently, the impact of different mycotoxins on cytokine secretion cannot be generalized across systems.

## 5. Conclusion

In summary, this study provides novel insight into how individual *Alternaria* mycotoxins differentially modulate the NF-κB signaling pathway in human THP-1 monocytes. By integrating functional readouts with transcriptional and protein-level analyses, as well as *in silico* docking analyses and downstream cytokine responses, distinct toxin-specific patterns of pathway interference were observed. The data suggest that these mycotoxins may affect NF-κB signaling at different regulatory levels, including upstream receptor signaling, IκBα regulation, kinase activity, and nuclear localization of NF-κB p65. In addition, receptor-mediated crosstalk, particularly involving GR-dependent mechanisms, may contribute to the observed immunomodulatory effects. Although all investigated mycotoxins suppressed NF-κB activity, the effects on individual pathway components differed considerably depending on the specific compound. This suggests that different *Alternaria* are likely to act through distinct mechanisms rather than a single common mode of inhibition. Collectively, these findings highlight the complex and multilayered nature of immunomodulation by *Alternaria* mycotoxins. The observed differences between individual compounds contribute to a better understanding of fungal toxin-host interactions and provide important information for future risk assessment, particularly regarding immune-related health effects.

## Abbreviations

AOH: Alternariol
AME: Alternariol monomethyl ether
ALTP: Alterperylenol
AST: Altersetin
ATX-I: Altertoxin I
AhR: Aryl-hydrocarbon receptor
CTB: CellTiter Blue
Dex: Dexamethasone
ELISA: Enzyme-linked immunosorbent
GR: Glucocorticoid receptor
IKK: IκB kinase
IκB: Inhibitor of κB
IL: Interleukin
LPS: Lipopolysaccharide
NF-κB: Nuclear factor kappa-light-chain-enhancer of activated B cells
qRT-PCR: Quantitative real time PCR
TLR4: Toll-like receptor 4
TNF-α: Tumor necrosis factor-alpha

## Acknowledgements

The authors thank the Core Facility Multimodal Imaging, member of the VLSI (Vienna Life Science Instruments), Faculty of Chemistry, University of Vienna, for technical assistance during microscopy-based workflows. This research was supported by the University of Vienna. The illustration in Figure 7 was generated using BioRender (https://BioRender.com/5fjkzh9).

## Data availability

The datasets generated during and/or analyzed during the current study are available from the corresponding author on reasonable request.

## Declaration of generative AI and AI-assisted technologies in the manuscript preparation process

During the preparation of this work the authors used ChatGPT in order to refine the language and improve the readability of the manuscript. After using this tool, the authors reviewed and edited the content as needed and take full responsibility for the content of the published article.

## Author contributions

Conceptualization: Doris Marko, Francesco Crudo; Methodology: Vanessa Partsch, Francesco Crudo, Christian Schröder, Giorgia Del Favero; Formal analysis and investigation: Vanessa Partsch; Writing - original draft preparation: Vanessa Partsch; Writing - review and editing: Francesco Crudo, Christian Schröder, Giorgia Del Favero, Doris Marko; Funding acquisition: Doris Marko; Supervision: Francesco Crudo, Doris Marko.

