## Supplementary material for "Modulation of NF-κB signaling by *Alternaria* mycotoxins: *in vitro* and *in silico* insights into molecular mechanisms of immunosuppression in THP-1 monocytes"

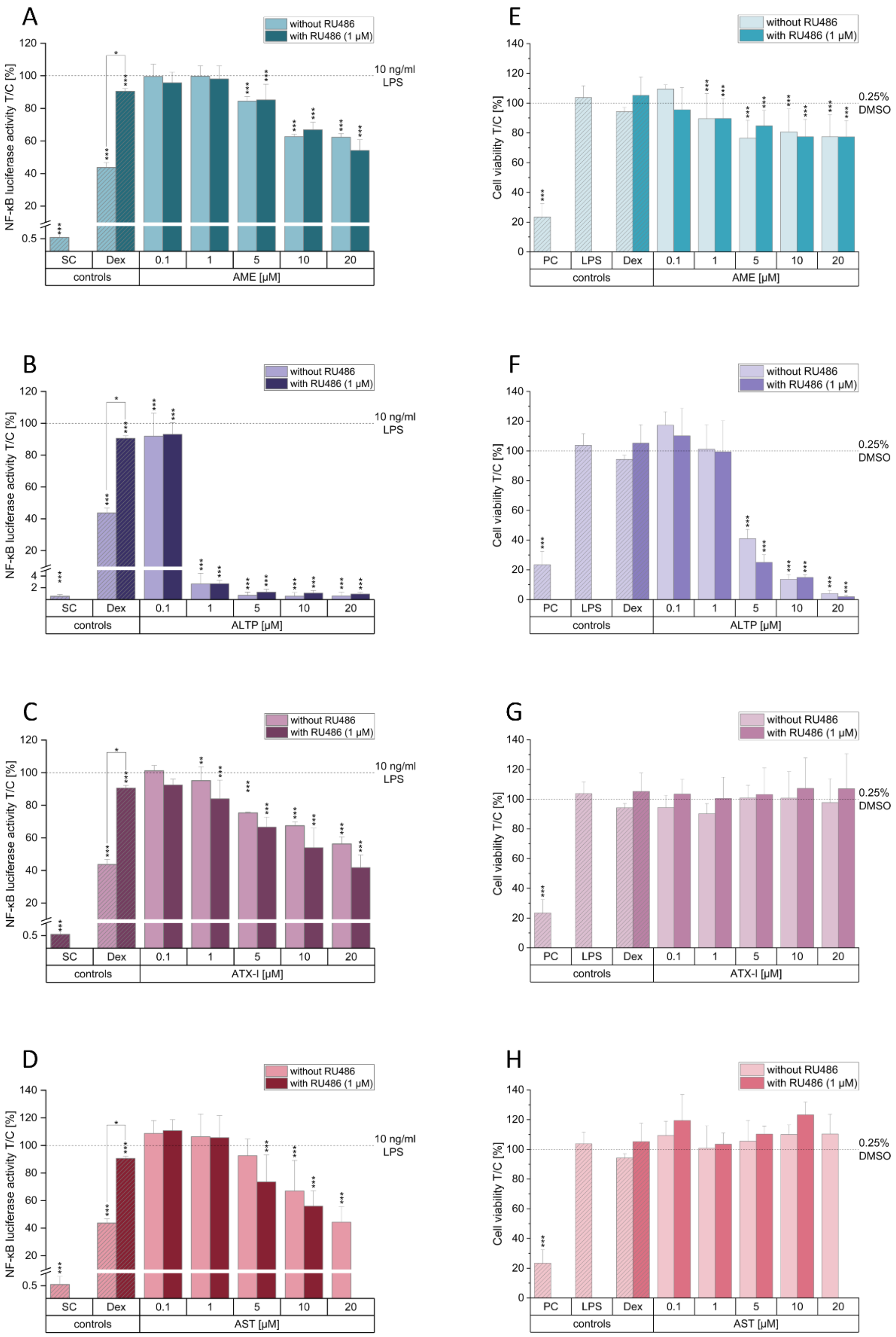


Supplementary Figure 1: Immunoinhibitory and cytotoxic effects of the *Alternaria* mycotoxins alternariol monomethyl ether (AME), alterperylenol (ALTP), altertoxin I (ATX-I) and altersetin (AST) in the presence and absence of the glucocorticoid receptor inhibitor RU486**.** THP-1 Lucia™ monocytes were either pre-incubated with 1 µM RU486 for 1 h or exposed to the *Alternaria* mycotoxins alone, followed by co-stimulation with 10 ng/mL lipopolysaccharide (LPS).Dexamethasone (Dex; 1 µM) served as negative control, 0.25% DMSO as the solvent control (SC), and in panels (E-H) 0.01% Triton X-100 as a positive control for cytotoxicity. Immunosuppressive effects were assessed by the NF-κB reporter gene assay (A-D) and cytotoxicity by the CellTiter-Blue® (CTB) assay (E-H). Data are presented as mean + SD of at least three independent experiments and are expressed relative to the positive control (10 ng/mL LPS) in (A-D) and relative to the SC in (E-H), indicated by dotted lines. Statistical significance compared to the positive control and between treatments with and without RU486 was evaluated using Student's *t*-test (*p<0.05, **p<0.01, and ***p<0.001).

Supplementary Table 1: Predicted docking of alternariol monomethyl ether (AME) to IκB kinase β (IKKβ) performed using SwissDock. Number of conformations per cluster and predicted binding free energies (ΔG) are shown. The lowest-energy conformation within each cluster was used for mapping the docking positions in Figure 8B.

| Cluster | Configurations | delta G | Position |
|---|---|---|---|
| 0 | 8 | -8.1 | 1 |
| 1 | 11 | -8 | 1 |
| 2 | 12 | -7.8 | 1 |
| 3 | 8 | -7.5 | 1 |
| 4 | 8 | -7.4 | 1 |
| 5 | 8 | -7.6 | 1 |
| 6 | 8 | -7.5 | 1 |
| 7 | 8 | -7.5 | 1 |
| 8 | 8 | -7.4 | 1 |
| 9 | 8 | -7.1 | 1 |
| 10 | 16 | -7.2 | 1 |
| 11 | 8 | -7.1 | 1 |
| 12 | 8 | -6.9 | 1 |
| 13 | 4 | -7.1 | 1 |
| 14 | 8 | -6.9 | 2 |
| 15 | 10 | -6.7 | 3 |
| 16 | 8 | -6.6 | 4 |
| 17 | 8 | -6.8 | 2 |
| 18 | 8 | -6.8 | 2 |
| 19 | 8 | -6.3 | 4 |
| 20 | 8 | -6.5 | 2 |
| 21 | 8 | -6.4 | 4 |
| 22 | 5 | -7.1 | 1 |
| 23 | 8 | -7 | 1 |
| 24 | 8 | -7 | 3 |
| 25 | 8 | -6.4 | 4 |
| 26 | 8 | -6.3 | 3 |
| 27 | 8 | -6 | 4 |
| 28 | 8 | -6.4 | 2 |
| 29 | 3 | -6.6 | 2 |
| 30 | 6 | -6.3 | 3 |
| 31 | 5 | -6 | 2 |

Supplementary Table 2: Predicted docking of alternariol (AOH) to IκB kinase β (IKKβ) performed using SwissDock. Number of conformations per cluster and predicted binding free energies (ΔG) are shown. The lowest-energy conformation within each cluster was used for mapping the docking positions in Figure 8C.

| Cluster | Configurations | delta G | Position |
|---|---|---|---|
| 0 | 3 | -8 | 1 |
| 1 | 8 | -7.8 | 1 |
| 2 | 10 | -7.7 | 1 |
| 3 | 8 | -7.7 | 1 |
| 4 | 8 | -7.6 | 1 |
| 5 | 8 | -8 | 1 |
| 6 | 8 | -7.7 | 1 |
| 7 | 8 | -7.5 | 1 |
| 8 | 8 | -7.5 | 1 |
| 9 | 8 | -7.4 | 1 |
| 10 | 8 | -7.5 | 1 |
| 11 | 8 | -7.2 | 1 |
| 12 | 3 | -7.7 | 1 |
| 13 | 8 | -7 | 1 |
| 14 | 5 | -7.1 | 2 |
| 15 | 9 | -7.3 | 1 |
| 16 | 8 | -6.7 | 1 |
| 17 | 16 | -6.5 | 3 |
| 18 | 6 | -6.5 | 2 |
| 19 | 8 | -6.4 | 3 |
| 20 | 8 | -6.5 | 4 |
| 21 | 8 | -6.5 | 3 |
| 22 | 5 | -6.9 | 1 |
| 23 | 8 | -6.5 | 2 |
| 24 | 3 | -6.7 | 2 |
| 25 | 8 | -6.6 | 4 |
| 26 | 7 | -6.5 | 4 |
| 27 | 4 | -6.6 | 5 |
| 28 | 8 | -6.6 | 4 |
| 29 | 8 | -6.2 | 2 |
| 30 | 8 | -6.2 | 2 |
| 31 | 2 | -6.1 | 2 |
| 32 | 8 | -6.5 | 5 |
| 33 | 8 | -6.3 | 6 |
| 34 | 1 | -5.8 | 5 |
| 35 | 3 | -5.4 | 5 |
| 36 | 1 | -5 | 4 |
| 37 | 2 | -2.1 | 1 |

Supplementary Table 3: Predicted docking of alterperylenol (ALTP) to IκB kinase β (IKKβ) performed using SwissDock. Number of conformations per cluster and predicted binding free energies (ΔG) are shown. The lowest-energy conformation within each cluster was used for mapping the docking positions in Figure 8C.

| Cluster | Configurations | delta G | Position |
|---|---|---|---|
| 0 | 8 | -8.4 | 1 |
| 1 | 6 | -8 | 1 |
| 2 | 16 | -7.7 | 1 |
| 3 | 8 | -7.6 | 2 |
| 4 | 8 | -8 | 1 |
| 5 | 8 | -7.3 | 1 |
| 6 | 8 | -7.5 | 1 |
| 7 | 8 | -7 | 3 |
| 8 | 8 | -7.4 | 4 |
| 9 | 8 | -6.6 | 5 |
| 10 | 24 | -6.3 | 5 |
| 11 | 8 | -6.3 | 5 |
| 12 | 8 | -7 | 2 |
| 13 | 1 | -7.6 | 1 |
| 14 | 8 | -6.8 | 5 |
| 15 | 8 | -7.9 | 1 |
| 16 | 3 | -6.7 | 3 |
| 17 | 8 | -6.6 | 3 |
| 18 | 8 | -6.4 | 6 |
| 19 | 8 | -7.1 | 7 |
| 20 | 8 | -7 | 1 |
| 21 | 8 | -6.9 | 2 |
| 22 | 5 | -6.7 | 3 |
| 23 | 8 | -7.2 | 8 |
| 24 | 8 | -6.1 | 4 |
| 25 | 8 | -6.9 | 4 |
| 26 | 8 | -6.2 | 4 |
| 27 | 8 | -6.9 | 1 |
| 28 | 8 | -6.2 | 5 |
| 29 | 8 | -6.3 | 5 |
| 30 | 8 | -6.7 | 4 |
| 31 | 1 | -6.2 | 1 |

Supplementary Table 4: Predicted docking of altertoxin I (ATX-I) to IκB kinase β (IKKβ) performed using SwissDock. Number of conformations per cluster and predicted binding free energies (ΔG) are shown. The lowest-energy conformation within each cluster was used for mapping the docking positions in Figure 8C.

| Cluster | Configurations | delta G | Position |
|---|---|---|---|
| 0 | 16 | -8.7 | 1 |
| 1 | 9 | -8.1 | 1 |
| 2 | 8 | -7.2 | 2 |
| 3 | 7 | -8 | 1 |
| 4 | 2 | -8.1 | 1 |
| 5 | 8 | -6.6 | 2 |
| 6 | 8 | -6.8 | 3 |
| 7 | 6 | -7.3 | 1 |
| 8 | 8 | -6.5 | 2 |
| 9 | 8 | -7.9 | 7 |
| 10 | 8 | -6.8 | 5 |
| 11 | 8 | -7 | 3 |
| 12 | 6 | -6.2 | 2 |
| 13 | 8 | -6.3 | 3 |
| 14 | 8 | -7 | 4 |
| 15 | 8 | -6.2 | 2 |
| 16 | 5 | -6.4 | 2 |
| 17 | 16 | -6.5 | 2 |
| 18 | 8 | -7.6 | 3 |
| 19 | 8 | -7.8 | 1 |
| 20 | 8 | -6.2 | 3 |
| 21 | 3 | -6.4 | 2 |
| 22 | 8 | -7 | 6 |
| 23 | 10 | -6.2 | 2 |
| 24 | 8 | -5.9 | 3 |
| 25 | 8 | -7 | 1 |
| 26 | 6 | -6.7 | 7 |
| 27 | 8 | -6.3 | 2 |
| 28 | 8 | -6.7 | 2 |
| 29 | 8 | -6.1 | 8 |
| 30 | 8 | -7.2 | 8 |
| 31 | 8 | -7.4 | 6 |
| 32 | 2 | -6.5 | 7 |

Supplementary Table 5: Predicted docking of altersetin (AST) to IκB kinase β (IKKβ) performed using SwissDock. Number of conformations per cluster and predicted binding free energies ($\Delta G$) are shown. The lowest-energy conformation within each cluster was used for mapping the docking positions in Figure 8C.

| Cluster | Configurations | delta G | Position |
|---|---|---|---|
| 0 | 3 | -7.3 | 1 |
| 1 | 8 | -7.9 | 2 |
| 2 | 8 | -7.5 | 1 |
| 3 | 8 | -7 | 3 |
| 4 | 8 | -7.4 | 1 |
| 5 | 8 | -6.9 | 3 |
| 6 | 8 | -7.1 | 1 |
| 7 | 16 | -7.3 | 3 |
| 8 | 4 | -6.7 | 4 |
| 9 | 8 | -7.1 | 3 |
| 10 | 8 | -7.7 | 2 |
| 11 | 3 | -6.3 | 3 |
| 12 | 8 | -7 | 1 |
| 13 | 8 | -7.4 | 2 |
| 14 | 8 | -7.5 | 1 |
| 15 | 8 | -7.3 | 1 |
| 16 | 8 | -7.2 | 1 |
| 17 | 8 | -6.7 | 3 |
| 18 | 8 | -6.5 | 3 |
| 19 | 8 | -6.4 | 3 |
| 20 | 8 | -6.8 | 1 |
| 21 | 8 | -6.9 | 2 |
| 22 | 8 | -6.9 | 3 |
| 23 | 8 | -6.8 | 3 |
| 24 | 8 | -6.8 | 3 |
| 25 | 5 | -6.8 | 3 |
| 26 | 4 | -6.2 | 4 |
| 27 | 8 | -6.9 | 1 |
| 28 | 8 | -6.7 | 5 |
| 29 | 8 | -6.8 | 3 |
| 30 | 5 | -6.5 | 1 |
| 31 | 5 | -7.7 | 2 |
| 32 | 8 | -6.6 | 3 |
| 33 | 8 | -6.9 | 1 |
| 34 | 3 | -7.6 | 2 |
